# A stepwise model for *asp* gene emergence across HIV and SIV lineages: progressive stop-codon loss and regulatory evolution

**DOI:** 10.64898/2026.09.03.749113

**Authors:** Miu Naruki, Asaka Yanada, Motofumi Saito, Shohei Nagata, Phillip K. Yamamoto, Hironori Sato, Akio Kanai

**Author notes:** Corresponding Author: Akio Kanai, PhD.

## Abstract

**Background:** The antisense protein (ASP), encoded by an open reading frame (ORF) overlapping the *env* gene in human immunodeficiency virus (HIV), was proposed over 30 years ago. ASP is an ∼189-amino-acid-long, highly hydrophobic protein with proposed roles in autophagy and viral entry. Although primarily described in pandemic HIV-1 group M, ASP was also identified in a single simian immunodeficiency virus (SIVcpz) isolate, suggesting recent acquisition. However, SIV from more distantly related lineages, including Old World monkeys (OWM), remains largely unexamined, and the broader origin of ASP is unresolved.

**Results:** We conducted computational analyses of 91,245 HIV-1, HIV-2, and SIV *env* sequences to assess the coding potential and evolutionary history of *asp* across primate lentiviruses. The sequences were classified into seven types based on the *asp*-frame ORF length and continuity: ASP, S1, and S2 for single continuous ORFs (≥ 450, 300‒449, and 120‒299 nucleotides, respectively); F1‒F3 for fragmented ORFs of comparable combined length; and Z for sequences lacking a detectable ORF of <120 nucleotides. Many SIV sequences retained a single, uninterrupted ORF of <120 nucleotides in the ASP frame, comparable to HIV-1 ASP-type sequences. A phylogenetic tree of 599 *env* sequences divided strains into HIV-1/SIVcpz/SIVgor and more distant HIV-2/SIV lineages. Fragmented *asp* ORFs were concentrated among basal, distantly related strains, suggesting *asp* emerged from stop-codon-rich ancestral segments. The stop-codon density progressively declined with ORF continuity in both groups (ρ = −0.63 and −0.77, respectively; both p < 0.001), consistent with the stop-codon-loss mechanism proposed previously for HIV-1 and extended to SIV here. The nonsynonymous/synonymous analysis revealed strong purifying selection within the overlapping region in HIV-1, and a more variable, lineage-dependent pattern in SIV. Motif analysis of the U3/LTR-like regions showed antisense promoter-associated transcription factor binding sites, including NF-κB and ETS1, were enriched in HIV-1, SIVcpz, and SIVgor.

**Conclusions:** These findings support a stepwise model of antisense regulatory evolution, whereby promoter-associated transcription factor motifs accumulated before a continuous *asp* ORF became fixed in HIV-1. Variation within the pandemic lineage suggests that the *asp* coding potential remains evolutionarily dynamic, providing new insights into the emergence and stabilization of overlapping genes during lentiviral evolution.

## Introduction

The emergence of new genes can provide viruses with adaptive advantages in replication, host interaction, immune evasion, pathogenesis, and transmission. Therefore, studying viral gene origins is central to understanding viral adaptation [1, 2]. New genes can arise through the reorganization of existing genetic material, including exon shuffling, gene duplication, retroposition, lateral gene transfer, and gene fusion or fission [3]. Alternatively, genes may arise *de novo* from non-coding regions or alternative reading frames within existing coding sequences, a process once considered rare but now recognized as an important source of evolutionary innovation [4]. Such genes frequently lack detectable homologs outside closely related lineages and may be classified as ORFans, although this absence alone does not establish a *de novo* origin [5, 6].

Viruses can expand their coding capacity through overprinting, in which a second open reading frame (ORF) emerges in an alternative reading frame within an existing protein-coding sequence, arising through point mutations that create a translation-initiation site, remove an interrupting stop codon, extend a pre-existing ORF, or a combination of these [1, 2, 6–8]. Because the overlapping proteins share the same nucleotide sequence, mutations can affect both reading frames simultaneously, imposing competing evolutionary constraints. Nevertheless, viruses have repeatedly used overprinting to expand their coding capacity without increasing genome size.

Human immunodeficiency virus (HIV) provides a useful system for investigating novel gene evolution because its error-prone replication generates high mutation rates and extensive genetic diversity, providing the sequence variation from which new coding regions may emerge and on which selection can act [9]. HIV-1 and HIV-2 arose through independent, cross-species transmissions of simian immunodeficiency viruses (SIVs) infecting African non-human primates. HIV-1 groups M and N were derived from SIVcpz in chimpanzees, groups O and P from SIVgor in gorillas, and HIV-2 from SIVsmm in sooty mangabeys [10–15]. In turn, SIVgor is thought to have originated from SIVcpz, which arose through recombination among SIV lineages infecting Old World monkeys (OWMs) [16]. Given this shared ancestry, HIV and SIV share a similar genome organization, comprising three core genes (*gag*, *pol*, and *env*), two essential regulatory genes (*tat* and *rev*), and several accessory genes (*vif*, *vpr*, *nef*, *vpx*, and *vpu*), whose presence and combination vary across lentiviral lineages. HIV-1 encodes nine canonical genes, although an ORF on the antisense strand, encoded in the −2 frame overlapping *env*, was proposed in 1988 as a 10^th^ gene encoding the putative *asp* gene (Supplementary Fig. S1) [17].

The *asp* ORF lies within the antisense transcript (AST) that is driven by a TATA-less negative-sense promoter in the U3 region of the 3′-LTR. This promoter is weaker than the positive-sense promoter, is indirectly repressed by Tat through promoter competition, and contains binding sites for specificity protein 1 (Sp1), nuclear factor kappa B (NF-κB), lymphoid enhancer-binding factor 1 (LEF1), ETS proto-oncogene 1 (ETS1), and upstream stimulatory factor (USF) [18–20]. AST is bifunctional, acting as an mRNA that produces ASP and, independently, as a regulatory long noncoding RNA to promote latency [21]. Antisense transcription is not unique to HIV-1; several deltaretroviruses encode conserved antisense proteins, whereas other retroviruses produce ASTs without detectable protein expression [21, 22]. This range, from transcript-only to protein-coding loci, shows that antisense transcription can exist without coding capacity, suggesting a regulatory framework may have preceded or accompanied the stabilization of ASP as a protein [21].

ASP is a highly hydrophobic protein, approximately 189 amino acids long, spanning the gp120/gp41 junction [17]. It is predicted to contain two transmembrane domains, two closely spaced cysteine triplets, and a proline-rich motif (PxxPxxP) resembling the SH3 domain-binding motif found in other proteins, including HIV-1 Nef, a motif that was recently implicated in ASP-mediated innate immune evasion [6, 23, 24]. Although the existence of a genuine ASP protein was long debated, several lines of evidence now support its expression, including the detection of *asp*-encoding transcripts and protein in HIV-1-infected cells and ASP-directed cellular immune responses comparable in frequency to those against Tat and Pol [25]. Proposed functions of ASP include autophagy induction, viral entry, and modulation of disease progression [6, 23, 26–28]. However, it remains difficult to study this protein owing to its low expression, hydrophobicity, and lack of clear homologs [25].

Despite increasing structural and functional characterization of ASP, its evolutionary origin and trajectory remain unresolved. A full-length *asp* ORF is associated predominantly with HIV-1 group M and is rare or absent in non-pandemic HIV-1 groups, HIV-2, and most examined SIV lineages. Only one SIVcpz sequence was reported to contain a comparable ORF [25]. This pattern was broadly corroborated by a recent narrative review that also identified *asp*-like antisense ORFs of variable lengths in selected SIVgsn, SIVmon, and SIVmus sequences [28]. Accordingly, a comprehensive comparison of HIV-1, HIV-2, and diverse SIV lineages is needed to determine whether shorter, fragmented, or positionally shifted antisense ORFs represent evolutionary precursors of *asp*, and how evolutionary constraint developed across the overlapping *env*/*asp* region.

Accessory genes in lentiviruses, which are often dispensable for replication, have repeatedly shaped host adaptation and, in some cases, pandemic emergence through their acquisition and refinement. We previously demonstrated that the HIV-1 accessory gene *vpu* originated in an ancestral SIV lineage infecting OWMs, rather than in SIVcpz directly, and that *vpu* underwent stepwise changes in its length, genomic organization, and functional specialization before becoming stabilized in HIV-1 group M [29]. These findings raise the possibility that *asp*, like *vpu*, emerged through a stepwise evolutionary process rather than through a single *de novo* event.

Here, we sought to address these questions by performing large-scale comparative analyses of 91,245 HIV-1, HIV-2, and SIV *env* sequences, classifying them into seven ASP types based on the ORF length and continuity as continuous ORFs (ASP, S1, and S2), fragmented ORFs (F1‒F3), and sequences lacking a detectable ORF of <120 nucleotides (nt) (Z). This framework allowed us to examine the start- and stop-codon distributions across the *asp*-like ORF structure, evaluate the evolutionary constraint within the *asp*-overlapping region by performing nonsynonymous (dN)/synonymous (dS) analysis using OLGenie, and search for U3/LTR-like regions for antisense-promoter-associated transcription factor (TF) motifs, including NF-κB and ETS1, using FIMO with dinucleotide-shuffled background controls. Together, these analyses tested whether the distribution of coding and regulatory features across primate lentiviruses is consistent with a stepwise model in which an antisense regulatory framework and partial coding potential preceded or accompanied the stabilization of a continuous *asp* ORF.

## Materials and Methods

### Datasets

The nucleotide and amino acid sequences of HIV and SIV strains used in this study were obtained from the National Center for Biotechnology Information (NCBI) GenBank (https://www.ncbi.nlm.nih.gov/, last accessed 16 July 2023) and the Los Alamos HIV Sequence Database (https://www.hiv.lanl.gov/, last accessed 23 July 2024), together with metadata on sampling region and year. In total, 100,920 *env* coding sequences (CDSs) were collected. Sequences were first filtered by length (>2,000 nt), yielding 91,218 sequences comprising 84,639 HIV-1 (groups M, N, O, and P), 49 HIV-2, and 6,530 SIV sequences spanning 28 lineages (Supplementary Table S1). This length-filtered set served as the starting dataset for ASP-type classification (see below) and, following additional redundancy reduction, phylogenetic analysis (see Molecular phylogenetic analysis).

### Multiple sequence alignment

The multiple sequence alignments of HIV and SIV of *env* nucleotide and amino acid sequences were conducted with MAFFT L-INS-i v7.45 [30] using the default parameters. The alignments of *env* nucleotide and amino acid sequences were visualized with Jalview v2.11 [31]. The percentage identity color scheme was applied with an 80% threshold, and the residues at alignment positions meeting or exceeding this threshold were colored pink to highlight the conserved regions. To quantitatively measure conservation, the numbers of amino acids with specific physicochemical properties conserved in each column of the alignment were calculated. This index was based on the Analysis of Multiply Aligned Sequences (AMAS) method [32], in which each alignment column receives a numerical conservation score reflecting the physicochemical similarity of the residues present, prioritizing identical residues followed by substitutions within the same physicochemical class. The conservation score for each amino acid position ranges from 1 to 11. High conservation is marked with * (score of 11 on the default amino acid grouping), whereas partial conservation in all properties is marked with a yellow + (score of 10) [32]. The consensus annotation shows the percentage of the modal residue per column. We used “+” to show that the modal value is shared by more than one residue. The alignments and tree files created in our study are provided in Supplementary Data 1 in the FASTA and Newick formats. The codon alignments for OLGenie analyses were generated by PAL2NAL using default parameters [33], which converts a multiple sequence alignment of proteins and their corresponding nucleotide sequences into a codon-based alignment. Specifically, the Env amino acid alignment created with MAFFT, together with the corresponding unaligned nucleotide sequences, was provided as the input into PAL2NAL. The resulting codon alignments were used to estimate the dS and dN substitution rates, and served as the input for all of the OLGenie analyses described below.

### ORF detection

The *asp* ORFs were first defined relative to the *env* region of HXB2 (GenBank K03455), the most widely used HIV-1 reference strain for functional studies. Following Cassan et al. 2016 [25], the HXB2 *asp* ORF is 189 codons long and located between *env* reference positions 1,717 and 1,151 in the −2 frame. Using these coordinates, we identified the corresponding *asp* region in other HIV and SIV *env* sequences by multiple sequence alignment against the *env* reference and examined the −2 frame for the *asp* start and in-frame stop codons. Classification was performed on 91,245 sequences in total, derived from the 91,218 sequences retained after initial filtering with two adjustments. First, for HIV-1 group O, only 17 of 88 complete *env* sequences passed the length-based filter. Because this lineage was underrepresented, all 88 complete *env* sequences were retained for ASP classification, subject to the *asp*-region alignment criteria described below. Second, 43 HIV-1 subtype B and C and 1 HIV-2 sequence that passed the initial length filter contained ambiguous bases, poor alignment, or other sequence irregularities that prevented reliable alignment to the *asp* query region and were therefore excluded. These adjustments yielded 91,245 sequences for ASP classification. GenBank accession numbers and sequence names are listed in Supplementary Data 2 according to their respective ASP type classifications. Numbers of sequences of each ASP type are provided in Supplementary Table S2. Only sequences that could be reliably aligned to the HXB2 *asp* region and contained both a start codon and a downstream in-frame stop codon were included in the classification.

The *asp* ORFs were visualized by aligning each sequence against a designated query, which was used to define the *asp* span together with a flanking window of ±200 nt in the alignment coordinates. The first start and first in-frame stop codons that aligned with the query within this window were recorded as putative *asp* ORFs. ORFs were only retained if both the start and stop positions fell entirely within the query-centered window. For each sequence, the overlapping ORFs were collapsed to the longest interval, and the alignment positions of the start and stop codons within the window were recorded as start/stop markers, thereby identifying *asp* segments within the reverse complement of all *env* sequences. The ORFs were subsequently extracted and translated for amino acid sequence alignments, and the sequences were categorized into types according to the ORF length and continuity. Start and stop codon counts within the *asp* window for the evolutionary group comparison (±150 nt flanking the *asp*-aligned region) were normalized per sequence and compared across ASP types using Dunn’s *post hoc* pairwise tests with *asp* as the reference group, and Cliff’s δ to quantify the effect size [34–36]. ASP types were treated as an ordered categorical variable representing increasing ORF length and continuity (Z → F3 → F2 → F1 → S2 → S1 → ASP). For each evolutionary group, Spearman’s rank correlation coefficients and associated p-values were calculated using scipy.stats.spearmanr (SciPy version 1.3.1) [37, 38] to test the monotonic association between ASP type rank and the per-sequence normalized start- or stop-codon rate. Tests were two-sided and conducted separately for each metric. Group 1 analyses included only the six observed categories from Z through S1.

### Molecular phylogenetic analysis

To reduce redundancy prior to phylogenetic tree construction, the sequences were clustered using CD-HIT at 80% identity [39] to group similar sequences and retain a single representative sequence per cluster. This step was applied to lineages with disproportionately large representation (HIV-1 subtypes A-G, HIV-2, SIVmac, SIVsab, and SIVsmm), producing a final set of 599 representative sequences used for phylogenetic analysis (Supplementary Table S1) ASP classification (described above) was performed on the final dataset of 91,245 sequences after the additional inclusion and exclusion steps, rather than on the CD-HIT-reduced set, to maintain adequate sample sizes across ASP types. All 48 HIV-2 sequences retained after these steps were included in the classification, despite being reduced to five representative sequences for the phylogenetic tree. The *env* nucleotide alignment of these 599 sequences (as described above) was trimmed using trimAl v1.2rev59, an automated tool for removing poorly aligned regions from sequence alignments. Applying the gappyout parameter, trimAl automatically detects and removes columns with excessive gaps, improving alignment quality; this trimming is particularly beneficial for largescale phylogenetic analyses [40]. An outgroup-rooted maximum likelihood tree, using SIVcol as the outgroup, was then constructed using IQ-TREE v1.6.12 [41]. We applied ultrafast bootstrap approximation (UFBoot2) [42], which enables rapid and unbiased estimation of branch support values, with 1000 bootstrap replicates. The best-fit substitution model was determined using ModelFinder [43]. We used the GTR+F+I+R10 model for Fig. 4, which combines the general time-reversible model, allowing variable substitution rates between nucleotides in both directions, with empirical base frequencies (F) and a FreeRate model with 10 rate categories to capture heterogeneity in evolutionary rate across sites [44–46].

### Estimating natural selection in the *asp* gene

Natural selection on the *asp* reading frame was quantified by dN/dS, specifically the dNN/dNS ratio for overlapping genes, where dNN represents substitutions that are nonsynonymous in both *env* and *asp*, and dNS represents substitutions that are nonsynonymous in *env* but synonymous in *asp*. Values of <1 indicate purifying selection, reflecting depletion of amino acid changing substitutions because of their likely deleterious effects, whereas values of ∼1 are consistent with neutral evolution, and values of >1 may indicate positive selection favoring amino acid changes. Because *asp* overlaps *env*, a reduced dN/dS in this region may reflect selective pressure on *env* and/or *asp*. Low dN/dS was therefore interpreted as evidence of constraint on the *asp*-overlapping region rather than direct proof of ASP protein function. Selection on the antisense *asp* ORF was quantified with OLGenie, an extension of dN/dS methods to overlapping genes [47]. The Env amino acid alignment and corresponding nucleotide sequences were converted to a codon alignment using PAL2NAL, which served as input for all dN/dS analyses. The sas12 overlapping frame (sense-antisense, with codon position 1 in *env* overlapping codon position 2 in the antisense *asp* ORF) was used, consistent with the known *env*/*asp* arrangement. The sites and pairwise differences were partitioned by OLGenie according to their effects in *env* and/or *asp*, using dN/dS as the primary measure of purifying selection on *asp*. Uncertainty was assessed via 1,000 nonparametric bootstrap replicates over the codon sites. The variation in selection along the overlapping region was examined using sliding windows of 25 codons, advanced one codon at a time across *env*. Windows were only retained if sufficient informative sites and at least six sequences were present, per OLGenie recommendations. The analyses were performed separately for HIV and SIV within each classified ASP type, excluding subsets of fewer than six sequences (e.g., SIV S1, containing only one sequence). The dN/dS values were compared between regions using the Mann‒Whitney *U* test, which was chosen because of its distribution-free assumptions and robustness for unequal group sizes, both of which are properties of sliding-window dN/dS data [48]. ΔdN/dS was assessed using percentile-bootstrap 95% confidence intervals (CIs) based on 10,000 resamples to quantify uncertainty in the mean difference without assuming a parametric sampling distribution [49]. The p-values in all statistical analyses were adjusted for multiple comparisons using the Holm‒Bonferroni method to control the probability of false positives under simultaneous testing [50].

### U3 region extraction and promoter motif analysis

For U3-region extraction, GenBank records corresponding to the type-specific FASTA entries were retrieved from the filtered sequence set summarized in Supplementary Table S1 using accession identifiers parsed from sequence headers. Because 3′-U3/LTR annotations were absent or inconsistently annotated across many GenBank records, candidate U3/LTR-like fragments were extracted using a BLAST anchoring approach [51, 52]. Sequences were grouped by evolutionary group, virus category, and ASP type, and the nucleotide records within each group were written to FASTA and indexed as BLAST databases using makeblastdb. To generate an initial anchor set, representative U3/LTR sequences from HIV-1 (K03455), SIVmac (M76764), and SIVcpz (OR117735) were used as query sequences. These queries were aligned against each GenBank-derived sequence set using blastn with -task blastn, word size 7, dust filtering disabled, and soft masking disabled. BLAST output was post-filtered to retain hits with ≥28% identity, alignment length of ≥40 nt, and an E-value of ≤10. For each GenBank record, the best-scoring hit was selected using bitscore. A sequence window extending 400 nt upstream and 400 nt downstream of the selected hit coordinates, bounded by sequence ends, was then extracted. Fragments were reverse complemented when the best hit aligned to the reverse strand so that the recovered regions were placed in a consistent orientation. To improve sensitivity across divergent HIV and SIV lineages, the candidate U3/LTR-like fragments recovered in the initial pass were pooled into an expanded multisequence query panel, which was used in the final extraction pass. Extracted fragments of ≥250 nt were retained as candidate U3/LTR-like sequences, and only fragments of 250‒800 nt were used to scan for TF binding motifs. Six TF binding motifs that were previously implicated in the HIV-1 antisense promoter (NF-κB, Sp1, LEF1, USF1, USF2, and ETS1) were retrieved from JASPAR (matrix ID in Supplementary Data 2) in the MEME format [53]. The occurrence of these motifs within each retained U3/LTR-like fragment was identified using FIMO in the MEME Suite [54]. The primary motif screen used a significance threshold of p ≤ 1 × 10^−3^, with an additional stricter threshold of p ≤ 1 × 10^−4^. Motif summaries were generated separately for matches in the positive and negative FIMO-reported motif orientations, as well as with both orientations combined. As a quality control step to reduce internal non-LTR BLAST matches, we evaluated whether the selected anchor hits were consistent with a 3′-LTR context based on their distance from the 3′ end of the GenBank record and, when available, nearby LTR/repeat-region or *nef* annotations. Motif summaries were then repeated after restricting the analysis to the 3′-LTR-consistent subset.

## Results and Discussion

### Identifying the ASP types according to the length and single or continuous ORFs

Although *asp* is known to be present in HIV-1 group M strains, its occurrence and evolutionary conservation in HIV-2 and SIV lineages other than SIVcpz/gor remain unclear. To systematically evaluate the *asp* coding potential across lentiviral lineages, we collected all available *env* sequences from the Los Alamos HIV Sequence Database, including 84,667 HIV-1 sequences from groups M, N, O, and P; 48 HIV-2 sequences; and 6530 SIV sequences from 28 SIV lineages. The putative *asp* ORFs were screened by aligning the reference *asp* query (reference) sequence with reverse-complemented *env* sequences, with start and stop codons in the −2 frame converted to uppercase to help identify the ORF relative to the query.

HIV-1 group M subtype A sequences have been reported to contain an early stop codon in *asp*, approximately 12 codons downstream of the annotated start, followed by an alternative downstream start codon approximately 17 codons later [25]. Consistent with this, the subtype A sequences in Fig. 1A contained a short upstream ORF fragment (∼33 nt) truncated by an early stop codon, followed by a second ATG initiating a longer downstream ORF (∼528 nt). Although no single continuous ORF spanned the full *asp* query region, the combined fragments totaled ∼561 nt, which is only ∼20 nt shorter than the reference *asp* query. Thus, subtype A may appear to lack *asp* under a strict continuous ORF definition while retaining substantial *asp* overlapping coding potential when considering the fragmented ORFs. To systematically examine this, we mapped the start codons, stop codons, and predicted ORFs in the *asp* frame within a 200-nt window flanking the query region (Fig. 1B), distinguishing continuous truncated ORFs. Based on this observation, we hypothesized that similar fragmented or downstream-shifted *asp* ORFs might also occur in other lentiviruses, including HIV-2 and SIVs, which remain largely underexplored. A recent review incorporating descriptive *in silico* analyses extended previous work to selected SIVgsn, SIVmon, and SIVmus sequences, and reported antisense ORFs of variable lengths retaining several *asp*-associated sequence features [28]. However, a systematic continuity-based comparison across the broader diversity of HIV-2 and SIV lineages remained unavailable. Because SIVcpz arose through recombination involving OWM SIV lineages, these more distantly related viruses may retain earlier stages of *asp*-like ORF evolution. To address this, we applied the same ORF-plotting approach to the broader HIV/SIV dataset, scanning the *asp*-aligned region and its flanking sequences (included because the exact *asp* position may vary among strains) for ATG-to-stop ORFs in the *asp* frame, and classifying each sequence by both ORF length and continuity. Using the full dataset, we observed two major patterns, sequences with one long continuous *asp* ORF, and sequences with fragmented *asp* ORFs, in which multiple shorter ORFs together overlapped the *asp* query region, similar to subtype A. Each sequence was categorized accordingly (Fig. 2; Supplementary Fig. S2). The ORF length distributions are shown in Supplementary Fig. S2 and representative ORF plots for each type in Supplementary Fig. S3. Sequences with a single continuous ORF were classified as ASP (≥450 nt, red), S1 (300‒450 nt, blue), or S2 (120‒299 nt, green). Sequences with multiple overlapping fragments were classified as F1 (combined length of ≥450 nt, orange), F2 (300‒449 nt, purple), or F3 (120‒299 nt, brown). Sequences with no detectable ORF of <120 nt were classified as Z type (grey). The sequence counts and average lengths for each type are shown in Fig. 2. In total, we classified 91,245 sequences, comprising 62,474 ASP, 363 S1, 6857 S2, 12,195 F1, 4658 F2, 2603 F3, and 2095 Z type.

**Fig. 1.**
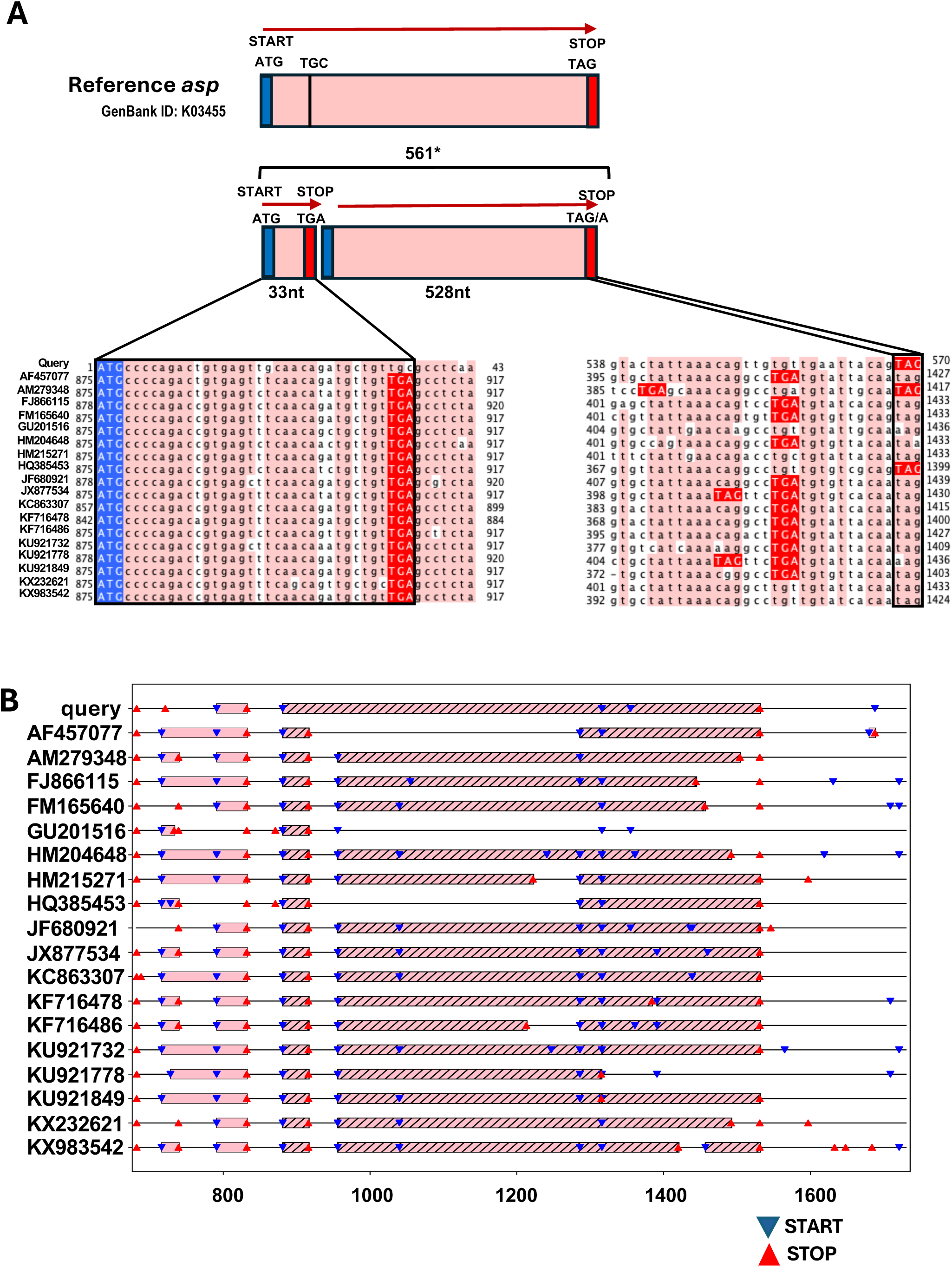
Identification of fragmented antisense protein (ASP). **A** Schematic overview and comparison of the reference *asp* and fragmented *asp*. HXB2, a widely used human immunodeficiency virus 1 (HIV-1) reference genome (GenBank ID: K03455), contains a single continuous *asp* open reading frame (ORF). In subtype A, the codon aligned to TGC in HXB2 is replaced by a premature stop codon (TGA), splitting the *asp*-overlapping region into a short upstream ORF and a longer downstream ORF. **B** ORF maps generated from the subtype A sequence alignment shown in (A), illustrating the resulting shortened and fragmented asp ORFs. Horizontal lines represent individual sequences; solid pink rectangles denote predicted −2 frame ORFs, and hatched segments indicate overlap with the reference asp ORF (K03455). Blue ▾ indicates start codons and red ▲ indicates stop codons. The shaded band marks the segment overlapping the reference asp ORF. Positions are indicated using alignment coordinates.

**Fig. 2.**
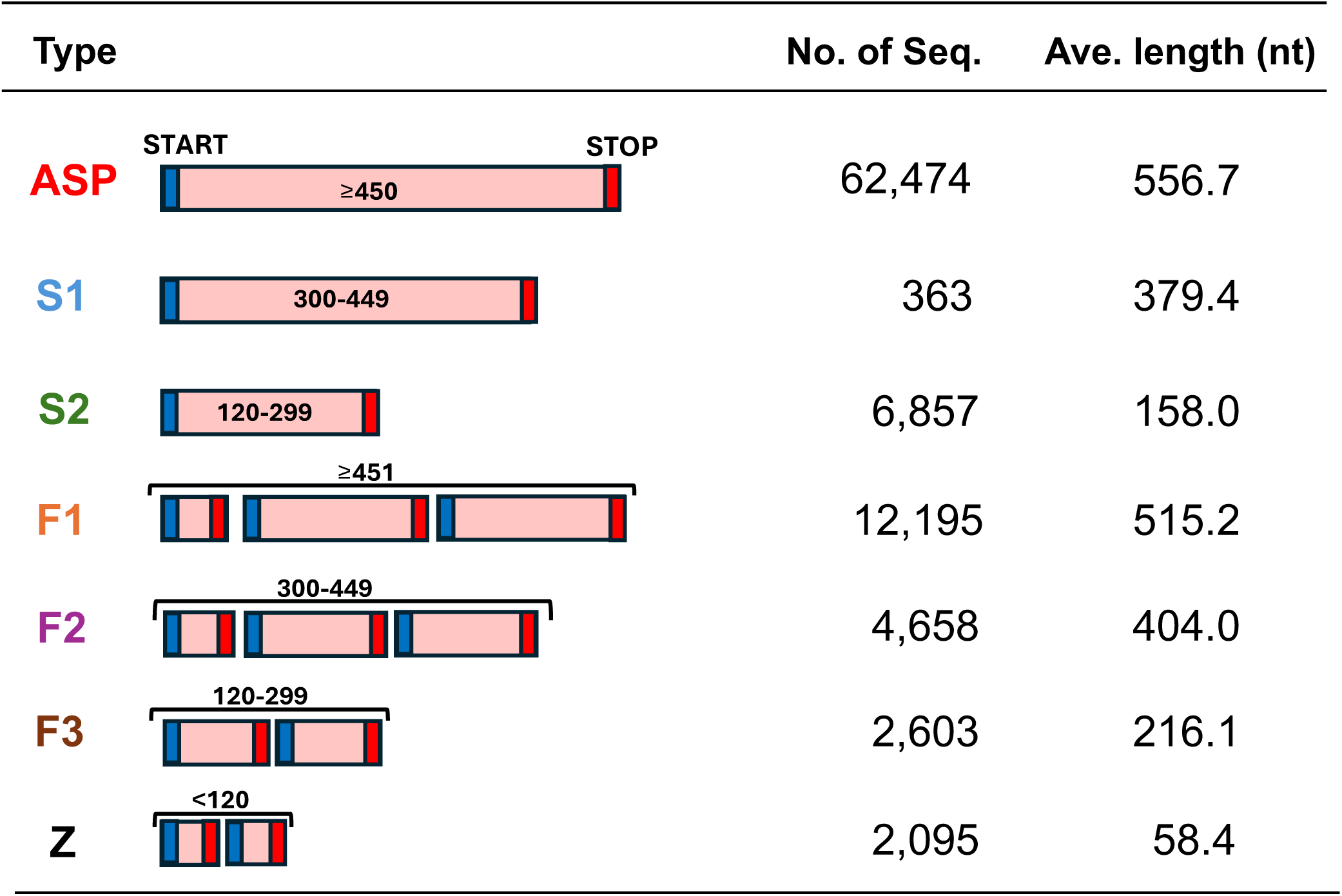
Classification of the ASP genes and their fragmented types used in this study. Full-length and fragmented ASP types were defined as follows: ASP (single continuous ORF ≥450 nt), S1 (300–449 nt), S2 (120–299 nt), and fragmented ASP types F1 (≥450 nt total), F2 (300– 449 nt total), F3 (120–299 nt total), and Z (<120 nt total). Blue bars indicate start codons (ATG), red bars indicate stop codons (TAA/TAG/TGA), and pink segments represent ORF regions. The number of sequences in each type and their average ORF lengths are shown next to each panel.

To evaluate the amino acid level conservation and compare differences among ASP types, we translated the representative sequences in the *env* −2 frame and performed multiple sequence alignment. Five representative sequences were selected from each of the ASP, S1, F1, and F2 types, allowing comparison of continuous and fragmented *asp* ORFs at the amino acid level (Fig. 3). In addition, we examined conservation within the ASP type by aligning 41 ASP-type amino acid sequences (Supplementary Fig. S4), which showed that conservation is concentrated in the N-terminal-to-mid region rather than being distributed across the full ORF. Conservation was strongest up to approximately amino acid 110, a region overlapping the Rev response element (RRE) on the positive strand, whereas the downstream region overlaps the hypervariable V5 (approximately residues 110‒130) and V4 (approximately residues 180‒200) regions of Env; conservation declined across this downstream region and was weakest in the final ∼30 C-terminal residues [55].

**Fig. 3.**
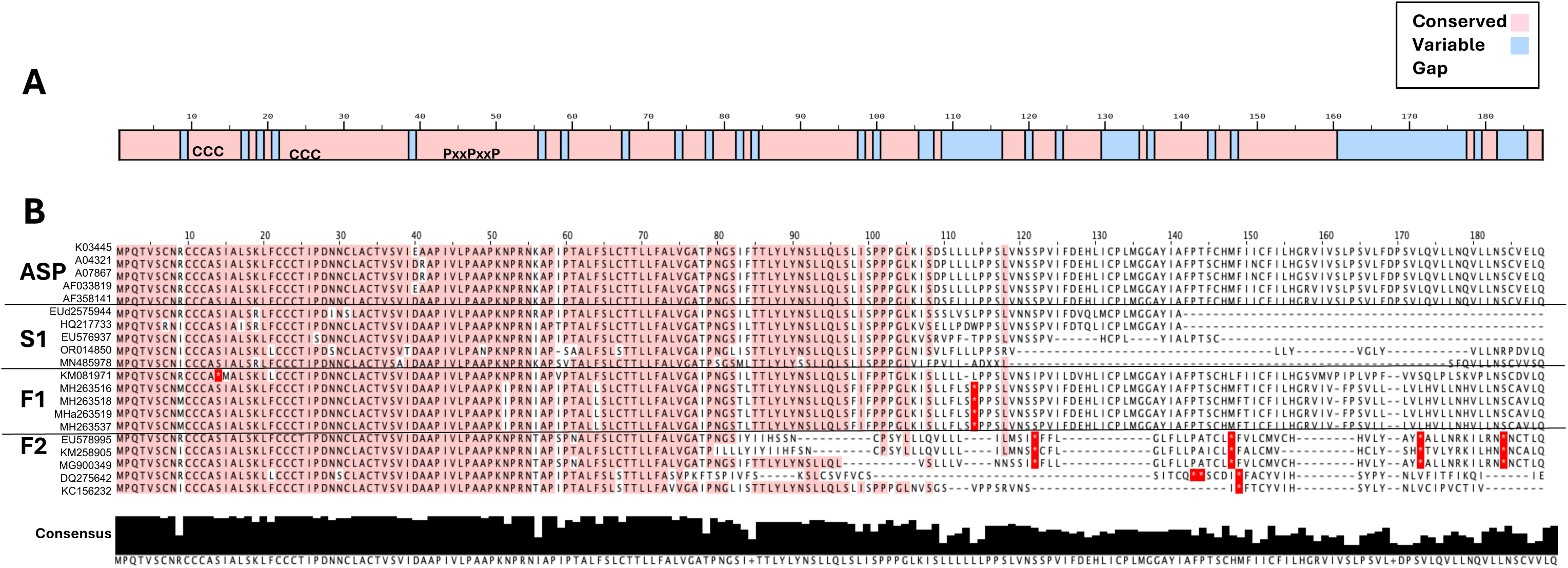
Summary of the amino acid sequence conservation in ASP and multiple sequence alignment of representative fragmented ASP types. **A** Summary of amino acid sequence conservation in *asp* genes. The conservation profile was generated from the *asp* gene alignment shown in Supplementary Fig. S4. Bars indicate the fraction of conserved residues (pink) and variable residues (blue) at each alignment position. **B** Multiple sequence alignment of ASP, S1, F1, and F2 types across the *asp* region. Residues with ≥80% identity are colored pink. Gaps are indicated using hyphens. Red * indicates stop codons. The consensus histogram shows the conserved regions. + indicates non-conserved residues.

Thus, ASP-type sequences retain a conserved core but not a uniformly conserved full-length sequence. To summarize the residue level conservation in continuous *asp* ORFs, we generated a conservation bar from the amino acid alignment of 41 ASP-type sequences as a reference profile for the *asp* region, confirming greater conservation across the N-terminal and central regions relative to the C-terminus (Fig. 3A). This profile was used to contextualize the representative ASP, S1, F1, and F2 alignment (Fig. 3B). The downstream region was more variable and contained more gaps and premature stop codons, especially in the fragmented types. Importantly, in the fragmented types, the residues downstream of premature stop codons should not be interpreted as part of a single continuous ASP protein because translation would terminate at the stop codon unless an alternative downstream start codon was used. Nevertheless, these downstream *asp*-frame regions often remained alignable with the reference ASP sequence and retained similar residues, particularly near the ASP C-terminus, possibly reflecting conservation of the overlapping *env* sequence on the opposite strand and/or preservation of downstream *asp* ORF fragments. This interpretation is consistent with previous reports indicating that the *asp* C-terminal intracellular region lacks obvious conserved motifs and is absent from some naturally truncated *asp* forms, suggesting it may be more dispensable than the N-terminal conserved motifs [6]. Notably, some previously described conserved motifs, namely the two cysteine-rich motifs (near alignment positions 10‒12 and 22‒24) and a PxxPxxP motif (at positions 43, 46, and 49) were retained across multiple classified types, suggesting that some *asp* sequence features are maintained even in shorter or fragmented forms. These cysteine-rich motifs have been proposed to form disulfide bonds or coordinate metal ions such as Zn^2^⁺ and Ni^2^⁺ [6]. However, the functional roles of these motifs in *asp* have not been experimentally confirmed. Overall, it seems that N-terminal *asp*-associated motifs are preferentially retained across ASP types, whereas the C-terminal region is more variable. Thus, further biochemical and functional studies are needed to determine whether these conserved motifs contribute to ASP function. Our findings extend recent reports of *asp*-like ORFs in selected OWM SIVs by placing these isolated observations within a broader continuum of fragmented, shortened, and continuous antisense coding potential.

### Loss of stop codons associated with the emergence of ASP in SIV and HIV

ORF plotting across the classified ASP types suggested that longer, continuous ORFs are associated with reduced stop-codon density in the *asp* frame, and the shorter and more fragmented types, particularly Z and F3, contained more stop codons within the *asp*-aligned region (Supplementary Fig. S3). To quantify this pattern, we counted the start (ATG) and stop (TAA/TAG/TGA) codons within the ASP window (the ASP-aligned region ±150 nt in alignment coordinates), and normalized the number per sequence (Supplementary Fig. S5). Dunn’s *post hoc* pairwise comparisons, using *asp* as the reference group and Cliff’s δ to summarize effect sizes, showed significantly fewer stop codons in the ASP-type sequences relative to all other types (δ = −0.14 to −0.89, indicating small to very large effects, with the largest differences in types with the most reduced ORF continuity). The start-codon density also differed significantly from those of *asp* (δ = +0.19 to +0.89), indicating generally greater abundance of start codons in *asp* sequences, albeit without the same clear progression across the types seen for stop codons. Ridgeline plots across the full *env* −2 frame alignment supported this distinction: the stop-codon density decreased progressively from the Z and fragmented types toward longer, continuous ASP ORFs, whereas the start-codon density showed a less consistent, type-dependent pattern (Supplementary Fig. S6). Subsequently, we checked whether the loss of stop codons was specific to the *asp* overlapping region or reflected a broader feature of the antisense-oriented *env* sequence. Across all ASP types, start and stop codons were less frequent within the *asp*-overlapping region than in the surrounding *env* −2 frame sequence, but the loss was more pronounced for stop codons (30‒38 vs. 43‒45 per 1,000 available codons outside the region), whereas the density of start codons showed less variability between regions. This indicates that the *asp*-overlapping region is relatively depleted of premature termination codons compared with the adjacent antisense *env* background. To examine whether ASP types were associated with phylogenetic relationships, we constructed a rooted phylogenetic tree using 599 *env* sequences from HIV-1, HIV-2, and SIV, using SIVcol as the outgroup (divergent and lacking a detectable *asp* ORF in our classification). The tree tips were colored by classified ASP type (Fig. 4). Based on the tree topology, we could divide the sequences into two evolutionary groups: Evolutionary Group 1 (HIV-2 and diverse SIV lineages other than SIVcpz/SIVgor) and Evolutionary Group 2 (HIV-1 groups M, N, O, P together with SIVcpz and SIVgor), reflecting the close evolutionary relationship among HIV-1, SIVcpz, and SIVgor. Within Evolutionary Group 2, the ridgeline plots of stop-codon density across the *env* −2 frame showed a clear reduction from the Z and fragmented types toward longer, continuous *asp* ORFs (Fig. 5; Supplementary Figs. S7–S9). A similar but less pronounced pattern was observed in Evolutionary Group 1 (Supplementary Fig. S7). Spearman’s rank correlation across the ordered ORF progression (Z → F3 → F2 → F1 → S2 → S1 → ASP) confirmed a significant negative association between the normalized stop-codon rate and ORF continuity in both groups (Group 1: ρ = −0.6314, n = 66, p < 0.001; Group 2: ρ = −0.7702, n = 529, p < 0.001). In that analysis, the correlation in Group 1 only spanned the six categories from Z through S1 because it contained no full-length ASP type sequences. However, this result does not imply that Evolutionary Group 1 lacks long antisense ORFs altogether because ORFs exceeding 150 codons were found in selected SIVmon and SIVmus genomes, for example [28]. The apparent difference between our finding of no full-length ASP-type sequences in Evolutionary Group 1 and this prior report of antisense ORFs exceeding 150 codons in SIVmon and SIVmus reflects the operational definitions used among studies; our definition of ASP type required a continuous ORF of ≥450 nt spanning the *asp* aligned region, whereas an ORF defined solely by its total length may begin from an alternative upstream start codon or differ in its positional correspondence to HIV-1 *asp*. Accordingly, Group 1 contained substantial partial ASP-like coding potential but no sequence that meets our definition of a full-length ASP type ORF. Representative alignments (Supplementary Figs. S7A and S8A) showed relatively high nucleotide conservation toward the *env* 3′ region corresponding to the *asp* N-terminus in both groups, with fragmented types retaining alignable, similar downstream sequences despite internal disruption. The corresponding ORF plots (Supplementary Figs. S7B and S8B) confirmed that the Z- and F3-type sequences contained numerous internal stop codons compared with the longer, more continuous types. We also examined whether the availability of a start codon tracked this transition. The ridgeline profiles of the start codon positions did not show a strong type- or group-dependent pattern (Supplementary Fig. S9), but the normalized per-sequence frequency of start codons was significantly and positively associated with the ordered ASP type in both groups (Group 1: ρ = 0.7086, n = 66, p < 0.001; Group 2: ρ = 0.7625, n = 529, p < 0.001). This suggests that the availability of start codons plays a secondary role in the *asp* ORF structure, but stop-codon loss remains the strongest and most consistent driver of ORF continuity. This finding is consistent with the results of Pavesi and Romerio [27], who proposed that the *asp* coding potential in HIV-1 is strongly influenced by the presence or absence of internal stop codons in the antisense frame. By extending this analysis to HIV-2 and diverse SIV lineages, we found that the association between stop-codon loss and *asp* ORF continuity is not restricted to pandemic HIV-1 group M. Both evolutionary groups showed reduced internal stop-codon rates toward more continuous ASP types, although the association was stronger and the type separation was more pronounced in Evolutionary Group 2, consistent with greater retention of *asp* coding potential within the HIV-1/SIVcpz/SIVgor clade. Together with the predominance of full-length ASP-type ORFs in the pandemic strain, this pattern raises the possibility that stabilization of the *asp* coding potential occurred during HIV-1 evolution and that ASP may have contributed to viral fitness in human hosts [25]. However, because this analysis does not fully account for phylogenetic non-independence and we did not reconstruct the ancestral states, we cannot confirm the direction or timing of this transition or distinguish progressive stabilization from independent gains and losses of ORF continuity. The presence of Z and fragmented types within HIV-1 further indicates that *asp* ORF continuity remains evolutionarily variable.

**Fig. 4.**
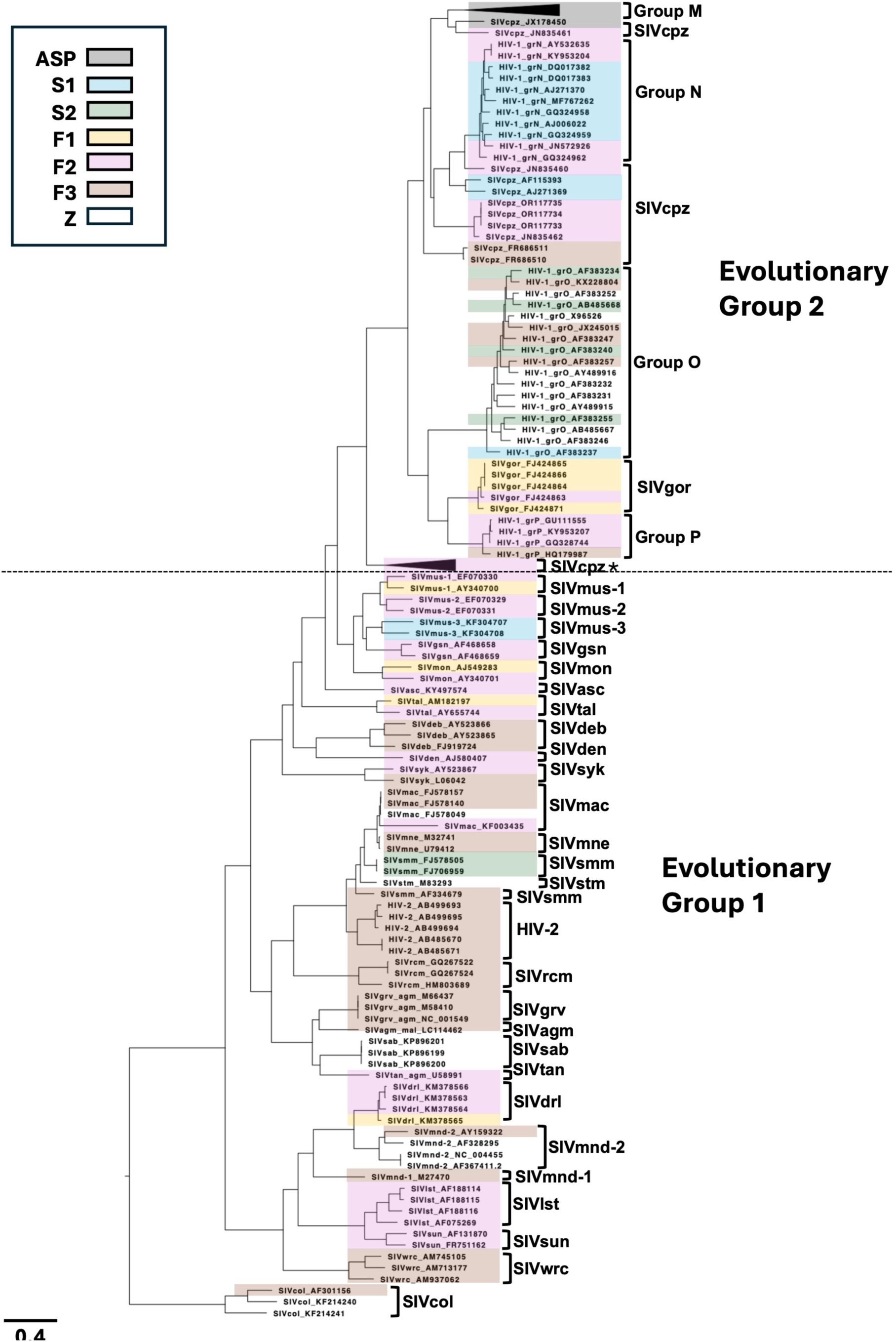
Rooted phylogenetic tree of *env* sequences. A rooted phylogenetic tree (rooted using SIVcol as the outgroup) was inferred from 599 nucleotide sequences with 1000 bootstrap replicates. The line splits Evolutionary Groups 1 and 2 in *asp* evolution. Tip annotations (viral strain and GenBank ID) are color-coded by ASP type: ASP = black; S1 = blue; S2 = green; F1 = yellow; F2 = pink; F3 = brown; Z = white. HIV-1 (group M) and SIVcpz clades are collapsed and shown as black ▲. The scale bar below the tree indicates 0.4 (40%) nucleotide substitutions per site.

**Fig. 5.**
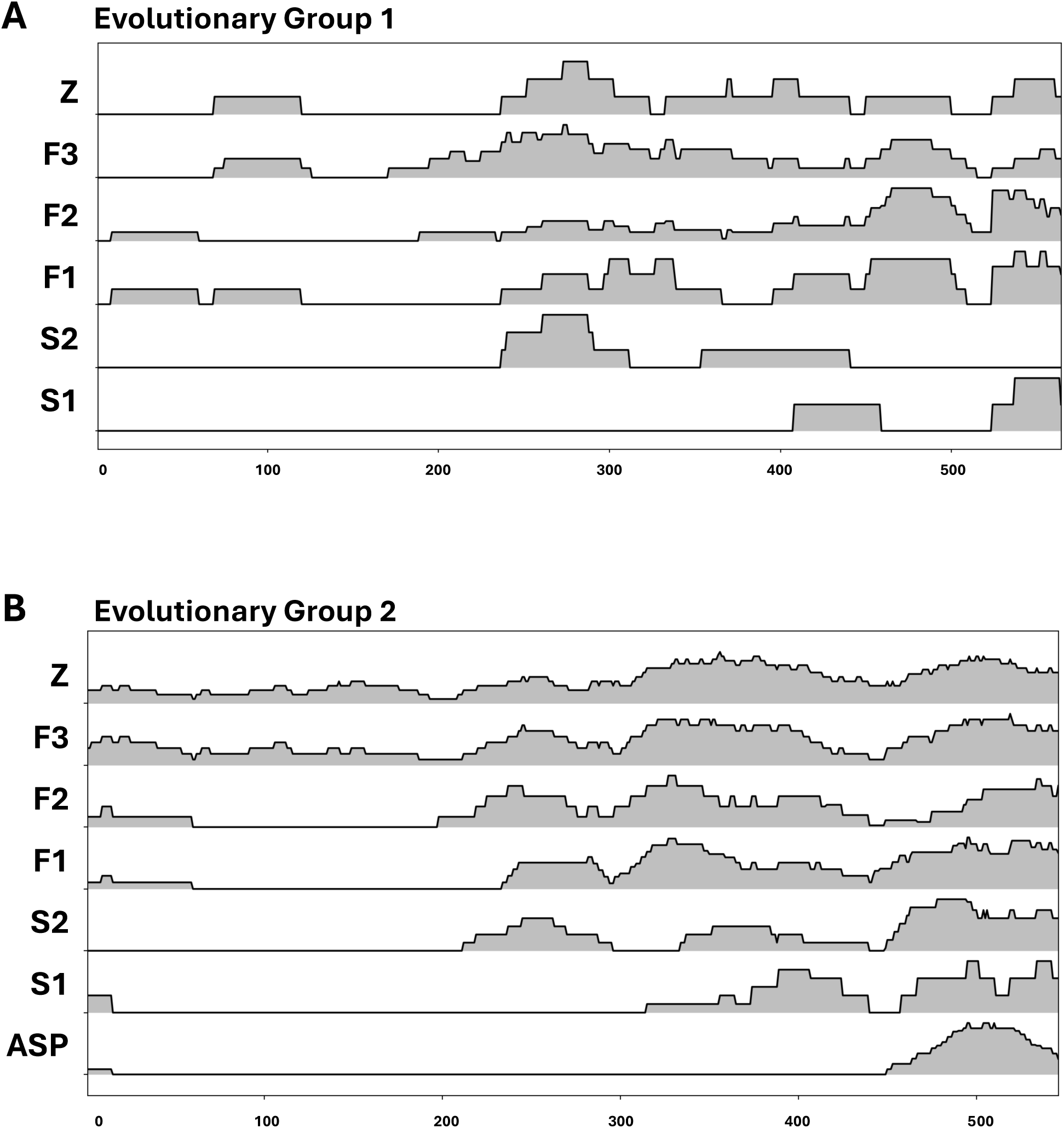
Evolutionary loss of stop codons underlying *asp* gene emergence in SIV and HIV within Evolutionary Groups 1. **(A) and 2 (B).** Ridgeline plots show the number of stop codons per type across the *asp* region in the 3′ → 5′ direction (query coordinates ± 100 nt; alignment coordinates). See Fig. 4 for reference.

### Evolutionary constraint and antisense promoter-like features across HIV and SIV

To assess the evolutionary constraint within the *asp*-aligned region and whether this differed between HIV-1 and SIV, we visualized dN/dS across the *env* coding sequence using sliding-window analysis for each ORF type (Supplementary Fig. S10). In the HIV-1 ASP type, dN/dS was consistently low within the *asp*-overlapping region, with values of ≤4 throughout much of the interval (Supplementary Fig. S10A). In SIV, the same region showed greater variability, with dN/dS values frequently approaching 1 or being >1, particularly toward the *asp* N-terminal side (Supplementary Fig. S10B). The Z type showed a comparatively constrained pattern in both HIV-1 and SIV, with dN/dS remaining largely <1 and nearing 0.4 in places (Supplementary Fig. S10C, D). SIV Z nevertheless showed a brief excursion to >1 toward the N-terminal side, representing an intermediate pattern between the stronger constraint observed in HIV-1 ASP and the more elevated values seen in other SIV types.

To quantify these patterns, we compared the mean dN/dS values between the *asp*-overlapping region and the flanking *env* sequence for each ORF type, computing ΔdN/dS as the difference between the overlap and flanking regions (i.e., overlap − flank; where negative values indicate lower dN/dS within the overlap), analyzing HIV-1 and SIV separately (Supplementary Table S3A). HIV-1 and SIV differed markedly. All seven HIV-1 ORF types showed significantly lower dN/dS values within the overlap than in the flanking sequence (all Holm‒Bonferroni-corrected p < 0.001; Cliff’s δ = −0.63 to −0.19). Averaged across the seven types, the mean dN/dS was 0.58 within the overlap and 0.73 outside the overlap (mean Δ = −0.15). This pattern was observed across the continuous, fragmented, and Z types, indicating that a reduction in dN/dS within the region was not restricted to sequences containing a single uninterrupted *asp* ORF. In contrast, SIV was more heterogeneous: S2 and F3 showed significantly higher dN/dS values within the overlap, F1 and F2 showed no significant difference, and Z showed significantly lower dN/dS values within the overlap, in the same direction as HIV-1. Averaged across the five analyzable SIV types, the mean dN/dS was 0.94 within the overlap and 0.89 outside the overlap (mean Δ = +0.06). Overall, HIV-1 showed a more consistent pattern of evolutionary constraint within the *asp*-overlapping region, whereas SIV exhibited a heterogeneous, ORF-type-dependent pattern. To further localize this lineage difference, we divided the *asp*-aligned interval into 5′ and 3′ halves, corresponding to the *asp* C-terminal and N-terminal regions, respectively, because *asp* is encoded antisense to *env* (Supplementary Table S3B). In HIV-1 ASP sequences, dN/dS was significantly lower in the N-terminal half than in the C-terminal half (0.417 versus 0.555, respectively; Δ[N-C] = −0.138, 95% CI −0.155 to −0.120, Holm-corrected p < 0.001, Cliff’s δ = −0.73), indicating stronger constraint toward the *asp* N-terminus. In contrast, most of the other HIV-1 ORF types (S2, F1, F3, and Z) showed significantly higher dN/dS values in the N-terminal half, and two (S1, F2) showed no significant difference between the C- and N-terminal halves. SIV showed a consistent pattern across all five analyzable types, with significantly higher dN/dS values in the N-terminal half in every case, indicating reduced constraint toward the *asp* N-terminal region. Thus, the pronounced N-terminal constraint observed in HIV-1 ASP sequences was not shared by most of the other HIV-1 ORF types or by the corresponding SIV groups. Notably, this same region, the *asp* N-terminus, overlaps the RRE on the positive strand, a cis-regulatory RNA element whose function depends on folding into a specific 3D shape rather than a fixed sequence. This structural, rather than sequence-level, constraint has been proposed to contribute to the region’s conservation [55]. The persistence of strong constraint in this region, specifically in HIV-1, is in contrast to its relaxation in SIV, and raises the possibility that the structural requirements of the RRE contribute an additional, HIV-1-specific layer of selective pressure superimposed on ASP and Env protein-coding constraint, one that may not apply equally across the SIV lineages examined here.

Because *asp* coding potential alone does not establish antisense transcription, we next investigated whether the recovered U3/LTR-like regions retained TF binding motifs associated with the HIV-1 NSP. We focused on six TF families previously implicated in this promoter (Sp1, NF-κB, ETS1, LEF1, USF1, and USF2) and scanned the U3/LTR-like fragments with FIMO. Using two reference genomes with annotated 3′-LTRs (HIV-1 MN685351, SIVmac M76764) to validate the workflow, all six families were recovered at p ≤ 1 × 10^−3^. At the stricter threshold of p ≤ 1 × 10^−4^, HIV-1 retained most signals while SIVmac retained a smaller core set (ETS1, Sp1, and NF-κB), confirming that the workflow could recover promoter-associated motifs from both reference sequences (Supplementary Fig. S11). Extending this to the full dataset (2568 HIV and 1408 SIV sequences), we compared the motif prevalence (fraction of sequences with ≥1 hit) against a dinucleotide-shuffled background (100 replicates per sequence) (Supplementary Table S4). This revealed clear lineage structure because NF-κB and ETS1 were depleted in Evolutionary Group 1 but enriched in Group 2, whereas USF1 tracked with the virus rather than lineage (i.e. depleted in HIV and enriched in SIV in both groups) and LEF1 was depleted across nearly all categories. Given that *asp* is antisense to the genomic reference, we then separately assessed enrichment by strand orientation within each virus/group combination (Supplementary Fig. S12A) and by individual ASP type (Supplementary Fig. S12B). The motif distributions varied by lineage, virus, and strand rather than being uniform. LEF1 was depleted in nearly all combinations, suggesting the selective loss of binding sites. NF-κB was enriched in Group 2 (both viruses and both strands) and depleted in Group 1 SIV (both strands), but was only depleted on the antisense strand in Group 1 HIV. ETS1 was broadly enriched, including in Group 2 SIV, but with strand-specific reversals being enriched only on the positive strand in Group 1 HIV, but depleted on the positive and enriched on the antisense strand in Group 2 HIV. This strand dependence is consistent with prior evidence showing that disrupting an ETS1 site reduces NSP activity even though ETS1 overexpression alone does not increase it, suggesting an indirect role that may involve cooperation with USF [20]. USF1 showed a virus-specific pattern that was independent of lineage or strand because it was depleted in HIV and enriched in SIV in all comparisons. This is consistent with prior work showing that USF sites are important for NSP activity, where even a single point mutation can abolish activity [18]. USF2 showed a similar pattern except in Group 2 SIV, where it was enriched on the positive strand but depleted on the antisense strand. Sp1 showed uniformly high raw prevalence (approximately 90%‒100% of sequences), but because shuffled controls also approached this ceiling, statistical enrichment was detected in only a subset of categories, a power limitation rather than evidence against a functional role. The ASP-type-level analysis clarified which classes drove these pooled patterns (Supplementary Fig. S12B). Sp1 and ETS1 were present in nearly every class (prevalence approaching 1.0). NF-κB was also present in most classes but dropped sharply in Group 1 SIV S2 (24.3%, n = 440) and Z (69.7%, n=412), accounting for the pooled Group 1 SIV depletion, while remaining close to 1.0 in F1 to F3. LEF1 fell to 3.6% in F3 (n = 445) but stayed high in S2 (93.6%) and Z (97.8%). USF2 showed the reverse pattern: high in F3 (82.5%), near 0 in S2 (3.9%) and Z (1.7%), and low across all Group 1 HIV classes (0% to 21.2%). This indicates that several pooled patterns were driven by specific classes rather than being lineage- or virus-wide; most notably, the depletion of NF-κB and LEF1 was particularly concentrated in Group 1 SIV types, whereas Sp1 and ETS1 remained near-universal. The results were robust at the stricter FIMO threshold (p ≤ 1 × 10^−4^); all six TF families were significantly enriched in the best-powered Group 2 classes (F1 and ASP), suggesting the reduced significance elsewhere reflects statistical power rather than the absence of a signal. These patterns broadly align with prior evidence implicating Sp1, NF-κB, and USF sites in HIV-1 antisense-promoter activity [18, 20]. The convergence of dN/dS, ORF continuity, and TF motif enrichment data is consistent with increased evolutionary constraint in the HIV-1 *asp*-overlapping region.

Collectively, these findings support a possible stepwise model of antisense regulatory evolution, in which promoter-associated TF motifs became enriched in the SIV lineage ancestral to HIV-1 (represented by SIVcpz and SIVgor) before the emergence and conservation of a continuous *asp* ORF. This order of events, where regulatory features appear before coding continuity, differs from more distantly related OWM SIV lineages, where these motifs were absent or were less frequent than in dinucleotide-shuffled control sequences. Our comparison captures a broad evolutionary transition rather than a precise timeline, so it does not directly show the order in which the individual features arose. Still, the overall pattern fits a model in which the regulatory framework and partial *asp* coding potential that was built up before a continuous reading frame became fixed in HIV-1, with gradual loss of internal stop codons possibly from fragmented to continuous *asp* ORFs.

## Acknowledgments

We thank members of the RNA Group at the Institute for Advanced Biosciences of Keio University, Japan, for insightful discussions.

## Author contributions

M.N. performed the investigation, data curation, formal analysis, and visualization and prepared the original draft of the manuscript. P.K.Y. contributed to script development and data analysis for the OLGenie analysis. P.K.Y., A.Y., M.S., S.N., and H.S. contributed to Writing – review & editing. A.K. conceived and supervised the study. M.N. and A.K. contributed to Writing – review & editing and finalized the manuscript. All authors reviewed and approved the final manuscript.

## Declarations

### Ethics approval and consent to participate

For this study, ethics approval and consent to participate were not required.

### Consent for publication

Not applicable.

### Availability of data and materials

The data supporting the findings of this work are available within the paper and its Supplementary Information files (Supplementary Data 1 and 2). The nucleotide and amino acid sequences of HIV and SIV strains used in this study were obtained from the National Center for Biotechnology Information (NCBI) GenBank (https://www.ncbi.nlm.nih.gov/, last accessed 16 July 2023) and the Los Alamos HIV Sequence Database (https://www.hiv.lanl.gov/, last accessed 23 July 2024), together with metadata on sampling region and year.

## Supplementary material

**Supplementary Table S1. Summary of *env* sequences used to construct the phylogenetic trees.**

The table lists complete *env* coding sequences (CDSs) collected from GenBank, the number of sequences that passed the minimum length filter (>2000 nt), the non-redundant set obtained after clustering with CD-HIT to remove nearly identical sequences (“CD-HIT”), and the final subset used to build the phylogenetic trees (“Tree”). *NA* not applicable.

**Supplementary Table S2. Distribution of nucleotide sequences used to construct the *asp* phylogenetic tree by type.**

Counts of sequences assigned to ASP, S1, S2, F1, F2, F3, and Z types that were included in the phylogenetic inference. For each type, refer to Fig. 2.

**Supplementary Table S3. Comparison of *env* selective constraint within and outside the *asp*-overlapping region across HIV and SIV ASP types.**

**A** *env* dN/dS values for the *asp*-overlapping region (inside *asp*) and the non-overlapping region (outside *asp*), shown for each ASP type in HIV-1 and SIV. ΔdN/dS was calculated as dN/dS (inside *asp*) − dN/dS (outside *asp*); negative values indicate lower dN/dS within the *asp*-overlapping region, consistent with stronger purifying constraint, whereas positive values indicate higher dN/dS within the ASP-overlapping region. % < 1 (inside *asp*) and % < 1 (outside *asp*) indicate the percentage of analyzed sites with dN/dS values of <1 within and outside the *asp*-overlapping region, respectively. **B** dN/dS in the 5′ and 3′ halves of the *asp*-overlapping region, by ORF type and virus. The *env* 5′ half and *env* 3′ half correspond to the *asp* C-terminus and N-terminus, respectively. Δ(5′−3′) is the 5′ half value minus the 3′ half value.

**Supplementary Table S4. Dataset-wide summary of transcription factor motif occurrence in HIV and SIV U3/LTR-like regions.**

Motif occurrence across the U3/LTR-like fragments extracted from HIV and SIV, grouped by evolutionary group and virus, using both-strand scanning at p ≤ 1 × 10^−3^. Columns display the number of sequences scanned (no. of seqs), the number with ≥1 hit (no. of seqs with hit), total motif matches (total hits), and the fraction of sequences with a hit (frac with hit), alongside the corresponding mean fraction across 100 dinucleotide-shuffled control replicates (frac with hit, shuffled control) and the empirical p-value (emp p-value) for that comparison. Asterisks denote significant enrichment and daggers indicate significant depletion relative to control (*/† p < 0.05, **/†† p < 0.01; ns = not significant).

**Supplementary Fig. S1. Schematic of the HIV-1 genome and the location of *asp*.** The *env* ORF is shown in the +1 frame (sense), and *asp* is located in the −2 frame (antisense), starting at its ATG initiation codon.

**Supplementary Fig. S2. Violin plots of the ORF length distributions for ASP types. A** Single, continuous ORFs: ASP (≥450 nt), S1 (300‒449 nt), and S2 (120‒299 nt). **B** Fragmented *asp* ORFs: F1 (≥450 nt), F2 (300‒449 nt), F3 (120‒299 nt), and Z (≤ <120 nt). The fragmented *asp* ORFs represent the distribution of the summed lengths of the individual ORFs.

**Supplementary Fig. S3. Visualization of the −2 frame ORF architectures in *env* across the classified ASP types.** Each row represents one sequence along the multiple alignment (*x*-axis: alignment position, nt). The query *asp* sequence is shown at the top. Horizontal lines indicate individual sequences and pink rectangles denote the predicted −2 frame ORFs. Hatched segments mark the overlap with the query-defined window. Blue▼indicates start codons and red▴indicates stop codons. Positions are shown in alignment coordinates, and the plotted window spans approximately 700–1700 nt (3′ → 5′ direction) on the alignment (≈ *asp* region ± 200 nt).

**Supplementary Fig. S4. Multiple sequence alignment and rooted phylogram of ASP amino acid sequences (ASP type).** The percentage identity color scheme was applied with an 80% threshold. Residues at alignment positions meeting or exceeding this threshold are colored pink to highlight the conserved regions. The histogram below the alignment shows physicochemical conservation, which also assigns high conservation to substitutions between amino acids with similar chemical properties, even when the residues are not identical. The phylogeny was inferred from the same amino acid alignment and rooted on the HIV-1 reference strain HXB2 (GenBank K03455) to orient the tree relative to a standard reference. The scale bar represents 0.09 amino acid substitutions per site.

**Supplementary Fig. S5. Boxplots comparing the start-and stop-codon counts within the *asp* window across ASP types.** The start-(ATG) and stop-(TAA/TAG/TGA) codon counts were computed within the strict *asp* region (no flanking sequence) and reported per sequence. **A** Start codons per sequence by type. **B** Stop codons per sequence by type. Boxes represent the interquartile range (IQR), the horizontal lines within the boxes show the median, whiskers extend to 1.5 × IQR, and dots indicate outliers. Each type was compared with *asp* (reference) using Dunn’s test; Holm-corrected p-values are shown across the six comparisons. The horizontal brackets above the plots show the significance and Cliff’s δ effect sizes. Significance: *** p < 0.001, ** p < 0.01, * p < 0.05; ns = not significant. Effect-size interpretation [1]: |δ| < 0.15 negligible, < 0.33 small, < 0.47 medium, ≥ 0.47 large.

1. Romano J, Kromrey JD. Appropriate statistics for ordinal level data: should we really be using t-test and cohen’s d for evaluating group differences on the nsse and other surveys? Paper presented at the Annual Meeting of the Florida Association of Institutional Research (FAIR).

**Supplementary Fig. S6. Ridgeline plots of the start and stop-codon counts in the *env* −2 frame. A** Start-codon (ATG) and **B** stop-codon (TAA/TAG/TGA) densities are shown as ridgeline plots for each type in the 3′ → 5′ direction. The *x*-axis represents alignment coordinates. The *y*-axis represents the counts normalized per sequence at each column (−2 frame only). The shaded band marks the *asp* region.

**Supplementary Fig. S7. Multiple sequence alignment and −2 frame stop-codon/ORF visualization for *asp* Evolutionary Group 1. A** Nucleotide multiple sequence alignment of representative Evolutionary Group 1 sequences. Positions sharing ≥80% identity with the *asp* query are highlighted in pink, and in-frame stop codons (TAA/TAG/TGA in −2 frame) are highlighted in red. **B** ORF map of the −2 frame across classes. Horizontal lines show the individual sequences, solid pink rectangles denote the predicted −2 frame ORFs, and hatched segments mark regions overlapping the query-defined window. Blue▼indicates start codons and red▴indicates stop codons. Positions are in alignment coordinates.

**Supplementary Fig. S8. Multiple sequence alignment and −2 frame stop-codon/ORF visualization for *asp* Evolutionary Group 2. A** Nucleotide multiple sequence alignment of representative Evolutionary Group 2 sequences. Positions sharing ≥80% identity with the *asp* query are highlighted in pink, and the in-frame stop codons (TAA/TAG/TGA in −2 frame) are highlighted in red. **B** ORF map of the −2 frame across classes. Horizontal lines show the individual sequences, solid pink rectangles denote the predicted −2 frame ORFs, and hatched segments mark the regions overlapping the query-defined window. Blue▼indicates start codons and red ▴ indicates stop codons. Positions are in alignment coordinates.

Supplementary Fig. S9. −2 Frame ORF mapping and start-codon ridgeline plots across the *asp* region in Evolutionary Groups 1 and 2. A schematic ORF maps of representative sequences from Evolutionary Groups 1 and 2 across the *asp* region. Horizontal lines represent the individual sequences, solid pink rectangles denote the predicted −2 frame ORFs, and hatched segments indicate the overlap with the query-defined window. Blue▼indicates start codons and red▴indicates stop codons. Positions are shown in alignment coordinates, spanning approximately 800‒1800 nt (the *asp* region ±200 nt flank). **B** ridgeline plots showing the per-type start-codon (ATG) density across the *asp* region, normalized per sequence. Positions are in query coordinates (±200 nt around the *asp* region).

**Supplementary Fig. S10. Sliding-window dN/dS across the *asp*-overlapping region in HIV-1 and SIV.** Sliding-window dN/dS profiles are shown: **A**, HIV-1 ASP type sequences; **B**, SIV sequences combined together; **C**, HIV-1 Z type sequences; and **D**, SIV Z type sequences. dN is the rate of nonsynonymous substitutions, whereas dS is the rate of synonymous substitutions. Accordingly, dN/dS values of <1 indicate purifying selection, values near 1 indicate neutral evolution, and values of >1 may indicate positive selection or relaxed constraint. The blue line shows the dN/dS values across codon-based sliding windows and the yellow band marks the hypothesized *asp*-overlapping region in alignment coordinates. Red dashed reference lines indicate dN/dS = 1.0, representing neutral evolution, and grey dashes dN/dS = 0.4, representing strong purifying constraint.

**Supplementary Fig. S11. Representative antisense promoter-associated TF motifs in HIV and SIV U3/LTR-like regions.** Schematic comparison of the canonical HIV-1 antisense promoter TF motif organization with representative HIV-1 (MN685351) and SIVmac (M76764) 3′-U3/LTR-like regions. The top schematic shows the typical arrangement of the TF binding motifs previously associated with HIV-1 antisense promoter activity. Representative HIV-1 and SIVmac LTR-like regions were extracted using GenBank LTR-related annotations or 3′-R-repeat landmarks and scanned with FIMO. Motifs shown for the representative sequences were retained at the stricter threshold of p ≤ 1 × 10^−4^.

**Supplementary Fig. S12. Distribution of predicted antisense TF promoter-associated motifs across HIV and SIV evolutionary groups at p ≤ 1 × 10^−3^. A** Grouped bar plots show strand-specific enrichment of each TF motif on the positive (+) and negative (−) strands, relative to a dinucleotide-shuffled sequence background (100 replicates per sequence). Each bar is one TF motif. The bar height is the difference between the observed and shuffled-background fraction hit (Δ = fraction of sequences with ≥1 FIMO hit, observed minus background), pooled across ASP types within each virus and evolutionary group. Positive bars indicate enrichment relative to the control and negative bars indicate depletion. **B** Heatmaps of the six TF motifs broken down by individual ASP type. Each cell depicts the fraction of sequences with ≥1 FIMO hits per TF motif (rows) in one ASP type. The panels are separated by virus and evolutionary group. Gray cells indicate ASP types with no available sequences in that category. In **A**, * denotes significant enrichment and † denotes significant depletion relative to the shuffled background (empirical p < 0.05, based on the distribution of 100 dinucleotide-shuffled replicates). Bars without a marker did not reach significance in either direction.

**Supplementary Data 1. Datasets and analysis outputs used to generate the figures presented in this article.**

**Supplementary Data 2. List of GenBank accession numbers by ASP type.**

