## Supplementary_tables for "A stepwise model for *asp* gene emergence across HIV and SIV lineages: progressive stop-codon loss and regulatory evolution"

*Author for Correspondence:

Akio Kanai, PhD

| **Strain** | | **Complete *env* CDS** | | **Filtered** | | **CD-HIT** | | **Tree** |
| --- | --- | --- | --- | --- | --- | --- | --- | --- |
| **HIV-1 A** |  | 954 |  | 951 |  | 18 |  | 18 |
| **HIV-1 B** |  | 55928 |  | 55741 |  | 98 |  | 98 |
| **HIV-1 C** |  | 26239 |  | 26218 |  | 90 |  | 90 |
| **HIV-1 D** |  | 1009 |  | 1008 |  | 12 |  | 12 |
| **HIV-1 F** |  | 184 |  | 184 |  | 11 |  | 11 |
| **HIV-1 G** |  | 473 |  | 473 |  | 18 |  | 18 |
| **HIV-1 H** |  | 5 |  | 5 |  | NA |  | 5 |
| **HIV-1 J** |  | 5 |  | 5 |  | NA |  | 5 |
| **HIV-1 K** |  | 2 |  | 2 |  | NA |  | 2 |
| **HIV-1 L** |  | 3 |  | 3 |  | NA |  | 3 |
| **HIV-1 VB** |  | 17 |  | 17 |  | NA |  | 17 |
| **HIV-1 N** |  | 11 |  | 11 |  | NA |  | 11 |
| **HIV-1 O** |  | 88 |  | 17 |  | NA |  | 17 |
| **HIV-1 P** |  | 4 |  | 4 |  | NA |  | 4 |
| **HIV-2** |  | 100 |  | 49 |  | NA |  | 5 |
| **SIVasc** |  | 1 |  | 1 |  | NA |  | 1 |
| **SIVcol** |  | 3 |  | 3 |  | NA |  | 3 |
| **SIVcpz** |  | 216 |  | 216 |  | NA |  | 216 |
| **SIVdeb** |  | 3 |  | 3 |  | NA |  | 3 |
| **SIVden** |  | 1 |  | 1 |  | NA |  | 1 |
| **SIVdrl** |  | 4 |  | 4 |  | NA |  | 4 |
| **SIVgor** |  | 5 |  | 5 |  | NA |  | 5 |
| **SIVgrv** |  | 3 |  | 3 |  | NA |  | 3 |
| **SIVgsn** |  | 2 |  | 2 |  | NA |  | 2 |
| **SIVlst** |  | 4 |  | 4 |  | NA |  | 4 |
| **SIVmac** |  | 314 |  | 285 |  | 5 |  | 5 |
| **SIVagm-mal** | | 1 |  | 1 |  | NA |  | 1 |
| **SIVmnd-1** |  | 1 |  | 1 |  | NA |  | 1 |
| **SIVmnd-2** |  | 4 |  | 4 |  | NA |  | 4 |
| **SIVmne** |  | 2 |  | 2 |  | NA |  | 2 |
| **SIVmon** |  | 2 |  | 2 |  | NA |  | 2 |
| **SIVmus-1** |  | 2 |  | 2 |  | NA |  | 2 |
| **SIVmus-2** |  | 2 |  | 2 |  | NA |  | 2 |
| **SIVmus-3** |  | 2 |  | 2 |  | NA |  | 2 |
| **SIVrcm** |  | 5 |  | 5 |  | NA |  | 3 |
| **SIVsab** |  | 125 |  | 125 |  | 3 |  | 3 |
| **SIVsmm** |  | 15185 |  | 5846 |  | 3 |  | 3 |
| **SIVstm** |  | 1 |  | 1 |  | NA |  | 1 |
| **SIVsun** |  | 2 |  | 2 |  | NA |  | 2 |
| **SIVsyk** |  | 2 |  | 2 |  | NA |  | 2 |
| **SIVtal** |  | 2 |  | 2 |  | NA |  | 2 |
| **SIVtan** |  | 1 |  | 1 |  | NA |  | 1 |
| **SIVwrc** |  | 3 |  | 3 |  | NA |  | 3 |
| **Total** |  | 100920 |  | 91218 |  | 599 |  | 599 |

**Supplementary Table S1. Summary of *env* sequences used to construct the phylogenetic trees.**

The table lists complete *env* coding sequences (CDSs) collected from GenBank, the number of sequences that passed the minimum length filter (>2000 nt), the non-redundant set obtained after clustering with CD-HIT to remove nearly identical sequences (“CD-HIT”), and the final subset used to build the phylogenetic trees (“Tree”). *NA* not applicable.

**Supplementary Table S2. Distribution of nucleotide sequences used to construct the *asp*** **phylogenetic tree by type.**

|  | | **ASP** | | **S1** | | **S2** | | **F1** | | **F2** | | **F3** | | | **Z** | |
| --- | --- | --- | --- | --- | --- | --- | --- | --- | --- | --- | --- | --- | --- | --- | --- | --- |
| **Total** | **Selected** | | **Total** | **Selected** | **Total** | **Selected** | **Total** | **Selected** | **Total** | **Selected** | **Total** | **Selected** | **Total** | **Selected** | | **Total** |
| **HIV-1 A** | 0 | | 7 | 0 | 0 | 0 | 4 | 14 | 693 | 1 | 169 | 2 | 63 | 1 | | 15 |
| **HIV-1 B** | 51 | | 43409 | 5 | 264 | 4 | 1014 | 10 | 7051 | 0 | 1580 | 6 | 1146 | 21 | | 1240 |
| **HIV-1 C** | 43 | | 17982 | 2 | 83 | 3 | 377 | 18 | 3994 | 4 | 2629 | 5 | 830 | 16 | | 318 |
| **HIV-1 D** | 6 | | 604 | 0 | 2 | 1 | 90 | 2 | 271 | 0 | 19 | 0 | 4 | 3 | | 18 |
| **HIV-1 F** | 5 | | 66 | 0 | 0 | 0 | 0 | 6 | 95 | 0 | 14 | 0 | 8 | 0 | | 1 |
| **HIV-1 G** | 17 | | 381 | 0 | 1 | 0 | 3 | 1 | 74 | 0 | 5 | 0 | 2 | 0 | | 7 |
| **HIV-1 H** | 1 | | 1 | 0 | 0 | 2 | 2 | 1 | 1 | 1 | 1 | 0 | 0 | 0 | | 0 |
| **HIV-1 J** | 3 | | 3 | 1 | 1 | 0 | 0 | 1 | 1 | 0 | 0 | 0 | 0 | 0 | | 0 |
| **HIV-1 K** | 1 | | 1 | 0 | 0 | 0 | 0 | 1 | 1 | 0 | 0 | 0 | 0 | 0 | | 0 |
| **HIV-1 L** | 2 | | 2 | 1 | 1 | 0 | 0 | 0 | 0 | 0 | 0 | 0 | 0 | 0 | | 0 |
| **HIV-1 VB** | 17 | | 17 | 0 | 0 | 0 | 0 | 0 | 0 | 0 | 0 | 0 | 0 | 0 | | 0 |
| **HIV-1 grN** | 0 | | 0 | 7 | 7 | 0 | 0 | 0 | 0 | 4 | 4 | 0 | 0 | 0 | | 0 |
| **HIV-1 grO** | 0 | | 0 | 0 | 0 | 4 | 16 | 0 | 2 | 1 | 11 | 4 | 32 | 8 | | 27 |
| **HIV-1 grP** | 0 | | 0 | 0 | 0 | 0 | 0 | 0 | 0 | 3 | 3 | 1 | 1 | 0 | | 0 |
| **HIV-2** | 0 | | 0 | 0 | 0 | 0 | 3 | 0 | 0 | 0 | 4 | 5 | 40 | 0 | | 1 |
| **SIVagm** | 0 | | 0 | 0 | 0 | 0 | 0 | 0 | 0 | 0 | 0 | 1 | 1 | 0 | | 0 |
| **SIVasc** | 0 | | 0 | 0 | 0 | 0 | 0 | 0 | 0 | 1 | 1 | 0 | 0 | 0 | | 0 |
| **SIVcol** | 0 | | 0 | 0 | 0 | 0 | 0 | 0 | 0 | 0 | 0 | 1 | 1 | 2 | | 2 |
| **SIVcpz** | 0 | | 0 | 2 | 2 | 1 | 1 | 4 | 4 | 202 | 202 | 5 | 5 | 2 | | 2 |
| **SIVdeb** | 0 | | 0 | 0 | 0 | 0 | 0 | 0 | 0 | 0 | 0 | 2 | 2 | 1 | | 1 |
| **SIVden** | 0 | | 0 | 0 | 0 | 0 | 0 | 0 | 0 | 1 | 1 | 0 | 0 | 0 | | 0 |
| **SIVdrl** | 0 | | 0 | 0 | 0 | 0 | 0 | 1 | 1 | 0 | 0 | 3 | 3 | 0 | | 0 |
| **SIVgor** | 0 | | 0 | 0 | 0 | 0 | 0 | 4 | 4 | 1 | 1 | 0 | 0 | 0 | | 0 |
| **SIVgrv** | 0 | | 0 | 0 | 0 | 0 | 0 | 0 | 0 | 0 | 0 | 3 | 3 | 0 | | 0 |
| **SIVgsn** | 0 | | 0 | 0 | 0 | 0 | 0 | 0 | 0 | 2 | 2 | 0 | 0 | 0 | | 0 |
| **SIVlst** | 0 | | 0 | 0 | 0 | 4 | 4 | 0 | 0 | 0 | 0 | 0 | 0 | 0 | | 0 |
| **SIVmac** | 1 | | 1 | 0 | 0 | 0 | 0 | 0 | 0 | 1 | 2 | 2 | 279 | 1 | | 3 |
| **SIVmnd-1** | 0 | | 0 | 0 | 0 | 0 | 0 | 0 | 0 | 0 | 0 | 1 | 1 | 0 | | 0 |
| **SIVmnd-2** | 0 | | 0 | 0 | 0 | 0 | 0 | 0 | 0 | 0 | 0 | 1 | 1 | 3 | | 3 |
| **SIVmne** | 0 | | 0 | 0 | 0 | 0 | 0 | 0 | 0 | 0 | 0 | 2 | 2 | 0 | | 0 |
| **SIVmon** | 0 | | 0 | 0 | 0 | 0 | 0 | 1 | 1 | 1 | 1 | 0 | 0 | 0 | | 0 |
| **SIVmus-1** | 0 | | 0 | 0 | 0 | 0 | 0 | 1 | 1 | 1 | 1 | 0 | 0 | 0 | | 0 |
| **SIVmus-2** | 0 | | 0 | 0 | 0 | 0 | 0 | 0 | 0 | 2 | 2 | 0 | 0 | 0 | | 0 |
| **SIVmus-3** | 0 | | 0 | 2 | 2 | 0 | 0 | 0 | 0 | 0 | 0 | 0 | 0 | 0 | | 0 |
| **SIVrcm** | 0 | | 0 | 0 | 0 | 0 | 0 | 0 | 0 | 0 | 0 | 3 | 5 | 0 | | 0 |
| **SIVsab** | 0 | | 0 | 0 | 0 | 0 | 0 | 0 | 0 | 0 | 0 | 0 | 0 | 3 | | 125 |
| **SIVsmm** | 0 | | 0 | 0 | 0 | 2 | 5343 | 0 | 0 | 0 | 1 | 1 | 170 | 0 | | 331 |
| **SIVstm** | 0 | | 0 | 0 | 0 | 0 | 0 | 0 | 0 | 0 | 0 | 0 | 0 | 1 | | 1 |
| **SIVsun** | 0 | | 0 | 0 | 0 | 0 | 0 | 0 | 0 | 2 | 2 | 0 | 0 | 0 | | 0 |
| **SIVsyk** | 0 | | 0 | 0 | 0 | 0 | 0 | 0 | 0 | 1 | 1 | 1 | 1 | 0 | | 0 |
| **SIVtal** | 0 | | 0 | 0 | 0 | 0 | 0 | 1 | 1 | 1 | 1 | 0 | 0 | 0 | | 0 |
| **SIVtan** | 0 | | 0 | 0 | 0 | 0 | 0 | 0 | 0 | 1 | 1 | 0 | 0 | 0 | | 0 |
| **SIVwrc** | 0 | | 0 | 0 | 0 | 0 | 0 | 0 | 0 | 0 | 0 | 3 | 3 | 0 | | 0 |
| **Total** | 147 | | 62474 | 20 | 363 | 21 | 6857 | 66 | 12195 | 231 | 4658 | 52 | 2603 | 62 | | 2095 |

Counts of sequences assigned to ASP, S1, S2, F1, F2, F3, and Z types that were included in the phylogenetic inference. For each type, refer to Fig. 2.

**Supplementary Table S3A. Comparison of *env* selective constraint within and outside the *asp*-overlapping region across HIV and SIV ASP types.**

| **ORF type** | **Virus** | **No. Codons (in *asp*)** | **No. Codons (outside *asp*)** | **Mean dN/dS (in *asp*)** | **Mean dN/dS (outside *asp*)** | **ΔdN/dS** | **95% CI** | **p (Holm-corrected)** | **Cliff's delta** | **Significant (alpha=0.05)** |
| --- | --- | --- | --- | --- | --- | --- | --- | --- | --- | --- |
| **ASP** | HIV | 400 | 1078 | 0.498 | 0.667 | -0.169 | [-0.187, -0.151] | <0.001 | -0.46 | Yes |
| **S1** | HIV | 188 | 621 | 0.549 | 0.846 | -0.297 | [-0.333, -0.261] | <0.001 | -0.63 | Yes |
| **S2** | HIV | 214 | 645 | 0.694 | 0.747 | -0.053 | [-0.095, -0.011] | <0.001 | -0.19 | Yes |
| **F1** | HIV | 296 | 753 | 0.585 | 0.655 | -0.070 | [-0.112, -0.025] | <0.001 | -0.25 | Yes |
| **F2** | HIV | 212 | 687 | 0.466 | 0.648 | -0.182 | [-0.215, -0.149] | <0.001 | -0.47 | Yes |
| **F3** | HIV | 223 | 660 | 0.618 | 0.781 | -0.163 | [-0.191, -0.135] | <0.001 | -0.43 | Yes |
| **Z** | HIV | 244 | 795 | 0.667 | 0.791 | -0.124 | [-0.144, -0.104] | <0.001 | -0.30 | Yes |
| **S2** | SIV | 180 | 603 | 1.022 | 0.907 | 0.115 | [0.051, 0.179] | <0.001 | 0.31 | Yes |
| **F1** | SIV | 175 | 597 | 0.934 | 0.853 | 0.081 | [0.027, 0.137] | 0.325 | -0.07 | No |
| **F2** | SIV | 181 | 638 | 0.965 | 0.859 | 0.106 | [0.043, 0.171] | 0.325 | -0.07 | No |
| **F3** | SIV | 180 | 633 | 0.941 | 0.866 | 0.075 | [0.038, 0.112] | <0.001 | 0.29 | Yes |
| **Z** | SIV | 168 | 611 | 0.855 | 0.939 | -0.084 | [-0.153, -0.010] | <0.001 | -0.28 | Yes |

**Supplementary Table S3B. Comparison of *env* selective constraint within and outside the *asp*-overlapping region across HIV and SIV ASP types.**

| **ORF type** | **Virus** | **No. codons (C-term)** | **No. Codon (N-term)** | **Mean dN/dS (C-term, *env* 5')** | **Mean dN/dS (N-term, *env* 3')** | **Δ dN/dS (N-term - C-term)** | **95% CI** | **p (Holm-corrected)** | **Cliff's delta** | **Significant (alpha=0.05)** |
| --- | --- | --- | --- | --- | --- | --- | --- | --- | --- | --- |
| **ASP** | HIV | 235 | 165 | 0.555 | 0.417 | -0.138 | [-0.155, -0.120] | <0.001 | -0.73 | Yes |
| **S1** | HIV | 88 | 100 | 0.513 | 0.580 | 0.066 | [0.016, 0.117] | 0.344 | 0.09 | No |
| **S2** | HIV | 100 | 114 | 0.531 | 0.836 | 0.306 | [0.246, 0.365] | <0.001 | 0.65 | Yes |
| **F1** | HIV | 138 | 158 | 0.488 | 0.669 | 0.180 | [0.114, 0.251] | 0.010 | 0.20 | Yes |
| **F2** | HIV | 92 | 120 | 0.459 | 0.471 | 0.013 | [-0.038, 0.064] | 0.344 | -0.11 | No |
| **F3** | HIV | 113 | 110 | 0.514 | 0.725 | 0.211 | [0.178, 0.246] | <0.001 | 0.77 | Yes |
| **Z** | HIV | 104 | 140 | 0.633 | 0.692 | 0.058 | [0.035, 0.082] | <0.001 | 0.32 | Yes |
| **S2** | SIV | 84 | 96 | 0.896 | 1.132 | 0.237 | [0.153, 0.318] | <0.001 | 0.46 | Yes |
| **F1** | SIV | 74 | 101 | 0.704 | 1.102 | 0.398 | [0.322, 0.477] | <0.001 | 0.61 | Yes |
| **F2** | SIV | 78 | 103 | 0.594 | 1.246 | 0.653 | [0.583, 0.721] | <0.001 | 0.97 | Yes |
| **F3** | SIV | 85 | 95 | 0.838 | 1.033 | 0.196 | [0.147, 0.245] | <0.001 | 0.49 | Yes |
| **Z** | SIV | 71 | 97 | 0.537 | 1.088 | 0.552 | [0.454, 0.649] | <0.001 | 0.65 | Yes |

**A** *env* dN/dS values for the *asp*-overlapping region (inside *asp*) and the non-overlapping region (outside *asp*), shown for each ASP type in HIV-1 and SIV. ΔdN/dS was calculated as dN/dS (inside *asp*) − dN/dS (outside *asp*); negative values indicate lower dN/dS within the *asp*-overlapping region, consistent with stronger purifying constraint, whereas positive values indicate higher dN/dS within the ASP-overlapping region. % < 1 (inside *asp*) and % < 1 (outside *asp*) indicate the percentage of analyzed sites with dN/dS values of <1 within and outside the *asp*-overlapping region, respectively. **B** dN/dS in the 5′ and 3′ halves of the *asp*-overlapping region, by ORF type and virus. The *env* 5′ half and *env* 3′ half correspond to the *asp* C-terminus and N-terminus, respectively. Δ(5′−3′) is the 5′ half value minus the 3′ half value.

| **Evolutionary group** | **virus** | **TF** | **No. of seqs** | **No. seqs with hit** | **total hits** | **frac with hit** | **frac with hit (shuffled)** | **emp p-value** | **significance** |
| --- | --- | --- | --- | --- | --- | --- | --- | --- | --- |
| **Group 1** | HIV | ETS1 | 165 | 165 | 928 | 1 | 0.937 | 0.0099 | ** |
| **Group 1** | HIV | NFKB | 165 | 165 | 1457 | 1 | 0.992 | 0.3069 | ns |
| **Group 1** | HIV | SP1 | 165 | 165 | 3351 | 1 | 1 | 0.9703 | † |
| **Group 1** | HIV | LEF1 | 165 | 110 | 217 | 0.667 | 0.901 | 1 | †† |
| **Group 1** | HIV | USF1 | 165 | 86 | 188 | 0.521 | 0.817 | 1 | †† |
| **Group 1** | HIV | USF2 | 165 | 31 | 79 | 0.188 | 0.728 | 1 | †† |
| **Group 1** | SIV | ETS1 | 1301 | 1301 | 6775 | 1 | 0.914 | 0.0099 | ** |
| **Group 1** | SIV | NFKB | 1301 | 817 | 3065 | 0.628 | 0.98 | 1 | †† |
| **Group 1** | SIV | SP1 | 1301 | 1301 | 30166 | 1 | 0.999 | 0.3267 | ns |
| **Group 1** | SIV | LEF1 | 1301 | 835 | 1670 | 0.642 | 0.91 | 1 | †† |
| **Group 1** | SIV | USF1 | 1301 | 1176 | 2946 | 0.904 | 0.836 | 0.0099 | ** |
| **Group 1** | SIV | USF2 | 1301 | 394 | 558 | 0.303 | 0.739 | 1 | †† |
| **Group 2** | HIV | ETS1 | 2403 | 2258 | 11045 | 0.94 | 0.901 | 0.0099 | ** |
| **Group 2** | HIV | NFKB | 2403 | 2400 | 49154 | 0.999 | 0.97 | 0.0099 | ** |
| **Group 2** | HIV | SP1 | 2403 | 2402 | 45916 | 1 | 0.999 | 0.3366 | ns |
| **Group 2** | HIV | LEF1 | 2403 | 2181 | 8419 | 0.908 | 0.926 | 1 | †† |
| **Group 2** | HIV | USF1 | 2403 | 1288 | 4903 | 0.536 | 0.764 | 1 | †† |
| **Group 2** | HIV | USF2 | 2403 | 1022 | 5729 | 0.425 | 0.668 | 1 | †† |
| **Group 2** | SIV | ETS1 | 107 | 107 | 280 | 1 | 0.847 | 0.0099 | ** |
| **Group 2** | SIV | NFKB | 107 | 107 | 2153 | 1 | 0.959 | 0.0396 | * |
| **Group 2** | SIV | SP1 | 107 | 107 | 1061 | 1 | 0.987 | 0.2079 | ns |
| **Group 2** | SIV | LEF1 | 107 | 17 | 44 | 0.159 | 0.954 | 1 | †† |
| **Group 2** | SIV | USF1 | 107 | 107 | 295 | 1 | 0.818 | 0.0099 | ** |
| **Group 2** | SIV | USF2 | 107 | 95 | 107 | 0.888 | 0.704 | 0.0099 | ** |

**Supplementary Table S4. Dataset-wide summary of transcription factor motif occurrence in HIV and SIV U3/LTR-like regions**

Dataset-wide summary of motif occurrence across extracted U3/LTR-like fragments from HIV and SIV, grouped by evolutionary group and virus, using both-strand scanning at p ≤ 1×10⁻³. Columns report the number of sequences scanned (No. of seqs), the number with at least one hit (No. seqs with hit), total motif matches (total hits), and the fraction of sequences with a hit (frac with hit), alongside the corresponding mean fraction across 100 dinucleotide-shuffled control replicates (frac with hit, shuffled control) and the empirical p-value (emp p-value) for that comparison. Asterisks denote significant enrichment and daggers significant depletion relative to control (* / † p<0.05, ** / †† p<0.01; ns = not significant)
