## Supplementary_figures for "A stepwise model for *asp* gene emergence across HIV and SIV lineages: progressive stop-codon loss and regulatory evolution"

***Author for Correspondence**:
Akio Kanai, PhD

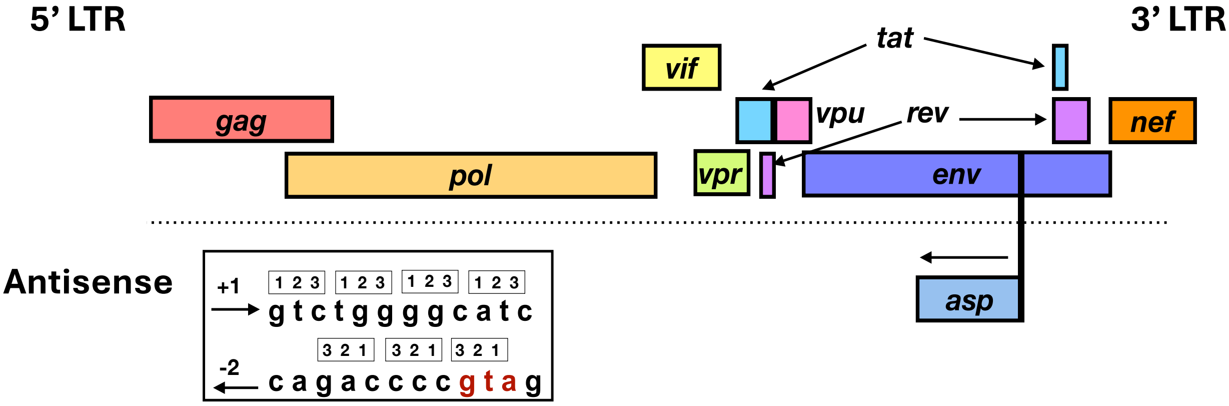


**Supplementary Fig. S1. Schematic of the HIV-1 genome and the location of *asp*.** The *env* ORF is shown in the +1 frame (sense), and *asp* is located in the −2 frame (antisense), starting at its ATG initiation codon.

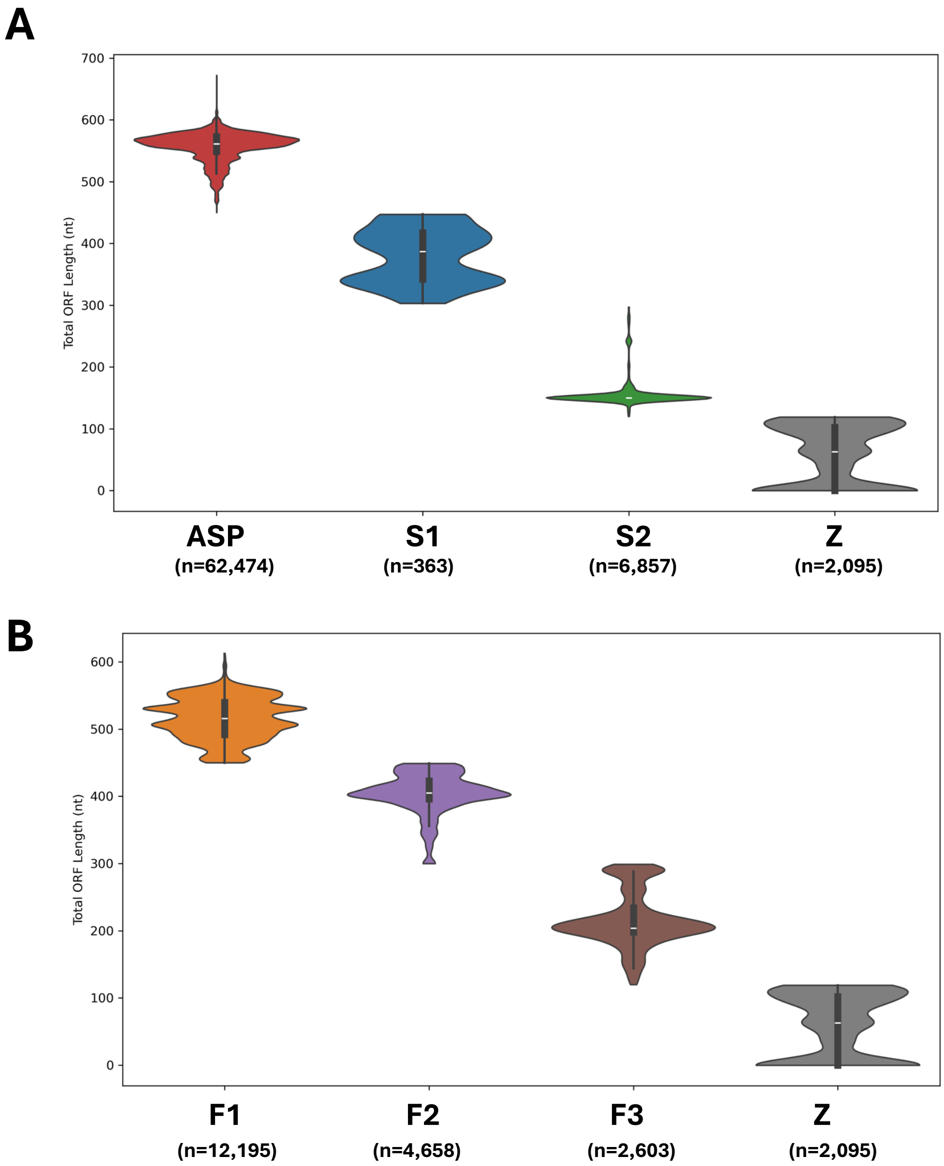


**Supplementary Fig. S2. Violin plots of the ORF length distributions for ASP types. A** Single, continuous ORFs: ASP (≥450 nt), S1 (300–449 nt), and S2 (120–299 nt). **B** Fragmented *asp* ORFs: F1 (≥450 nt), F2 (300–449 nt), F3 (120–299 nt), and Z (<120 nt). The fragmented *asp* ORFs represent the distribution of the summed lengths of the individual ORFs.

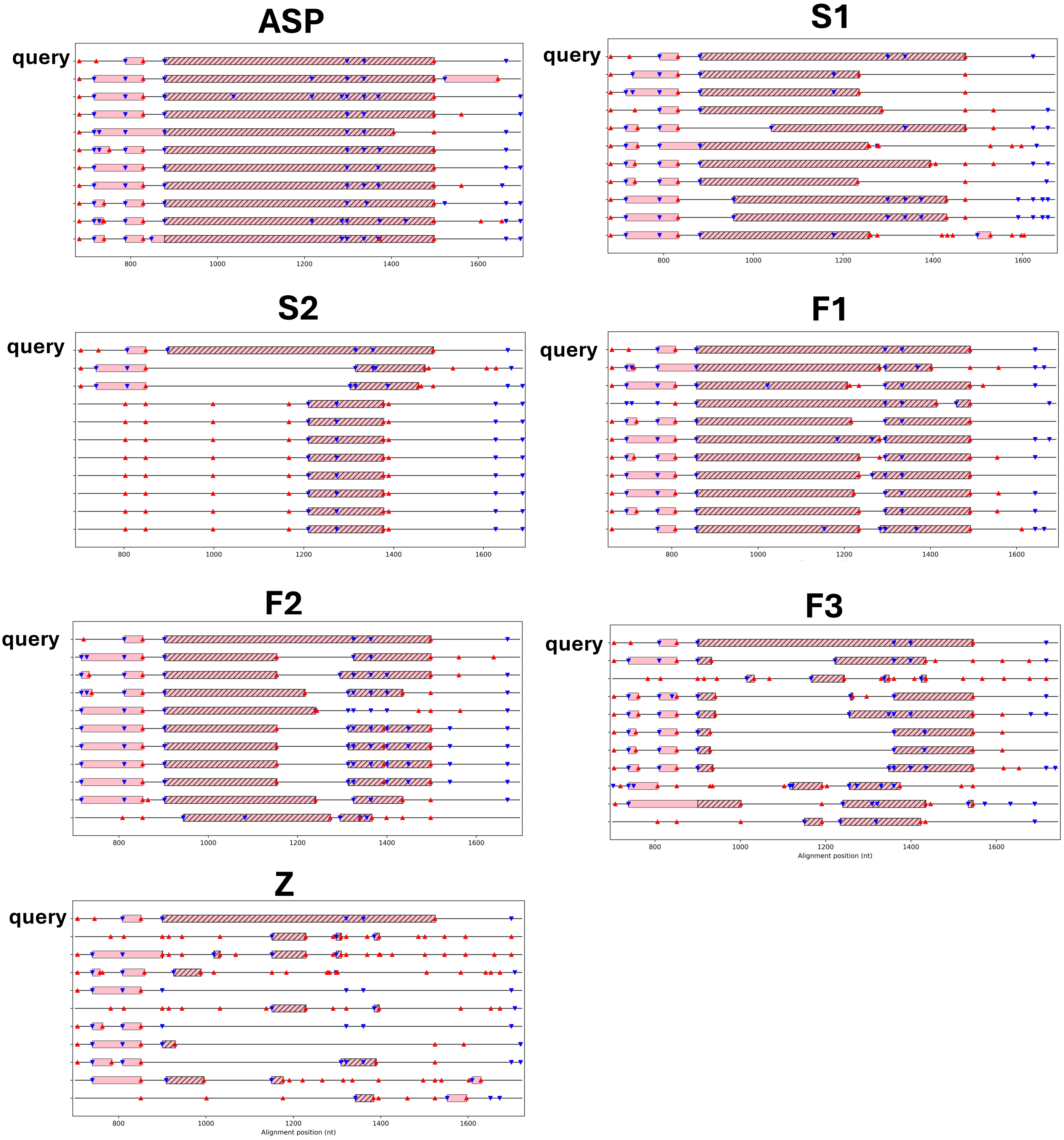


**Supplementary Fig. S3. Visualization of the −2 frame ORF architectures in *env* across the classified ASP types.** Each row represents one sequence along the multiple alignment (*x*-axis: alignment position, nt). The query *asp* sequence is shown at the top. Horizontal lines indicate individual sequences and pink rectangles denote the predicted −2 frame ORFs. Hatched segments mark the overlap with the query-defined window. Blue ▼ indicates start codons and red ▲ indicates stop codons. Positions are shown in alignment coordinates, and the plotted window spans approximately 700–1700 nt (3′ → 5′ direction) on the alignment (≈ *asp* region ± 200 nt).

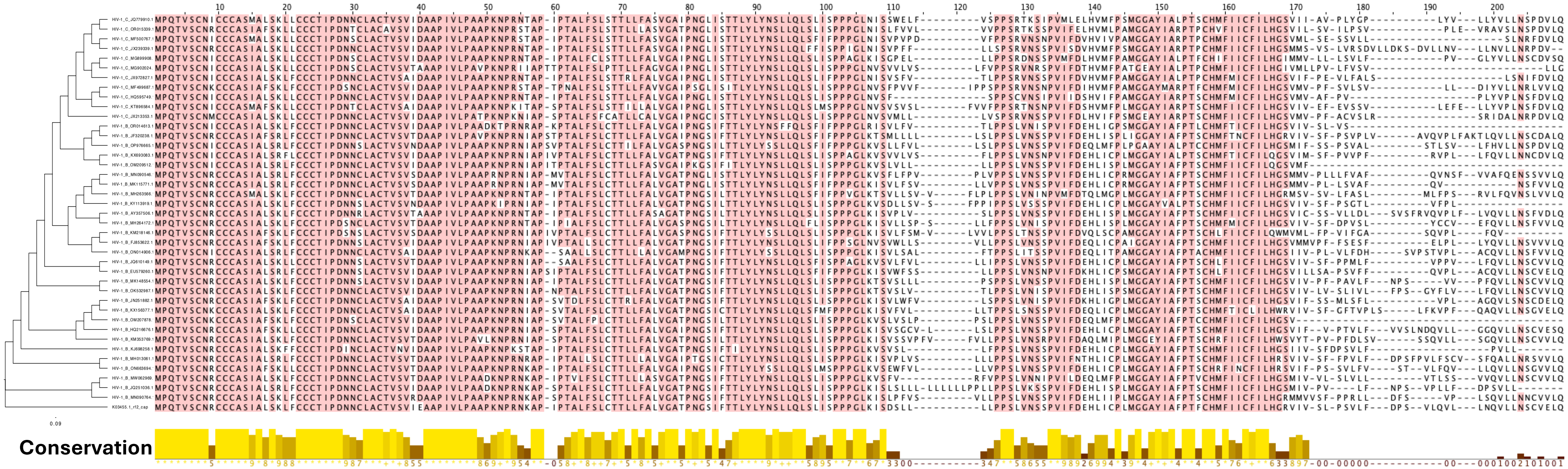

**Supplementary Fig. S4. Multiple sequence alignment and rooted phylogram of ASP amino acid sequences (ASP type).** The percentage identity color scheme was applied with an 80% threshold. Residues at alignment positions meeting or exceeding this threshold are colored pink to highlight the conserved regions. The histogram below the alignment shows physicochemical conservation, which also assigns high conservation to substitutions between amino acids with similar chemical properties, even when the residues are not identical. The phylogeny was inferred from the same amino acid alignment and rooted on the HIV-1 reference strain HXB2 (GenBank K03455) to orient the tree relative to a standard reference. The scale bar represents 0.09 amino acid substitutions per site.

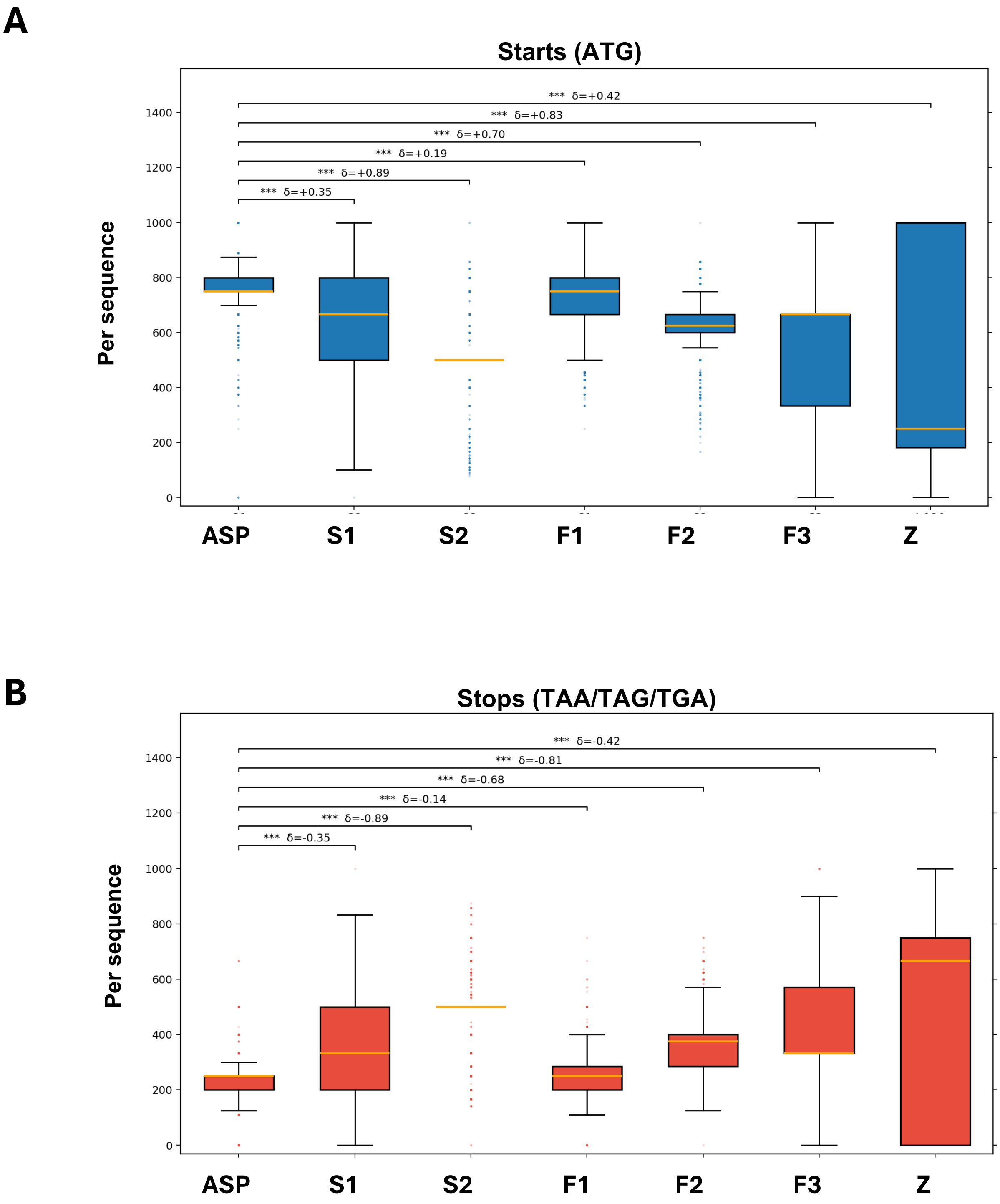


**Supplementary Fig. S5. Boxplots comparing the start- and stop-codon counts within the *asp* window across ASP types.** The start- (ATG) and stop- (TAA/TAG/TGA) codon counts were computed within the strict *asp* region (no flanking sequence) and reported per sequence. **A** Start codons per sequence by type. **B** Stop codons per sequence by type. Boxes represent the interquartile range (IQR), the horizontal lines within the boxes show the median, whiskers extend to 1.5 × IQR, and dots indicate outliers. Each type was compared with *asp* (reference) using Dunn's test; Holm-corrected p-values are shown across the six comparisons. The horizontal brackets above the plots show the significance and Cliff's δ effect sizes. Significance: *** $p < 0.001$, ** $p < 0.01$, * $p < 0.05$; ns = not significant. Effect-size interpretation [1]: $|\delta| < 0.15$ negligible, $< 0.33$ small, $< 0.47$ medium, $\geq 0.47$ large.

1. Romano J, Kromrey JD. Appropriate statistics for ordinal level data: should we really be using t-test and cohen's d for evaluating group differences on the nsse and other surveys? Paper presented at the Annual Meeting of the Florida Association of Institutional Research (FAIR).

**A**

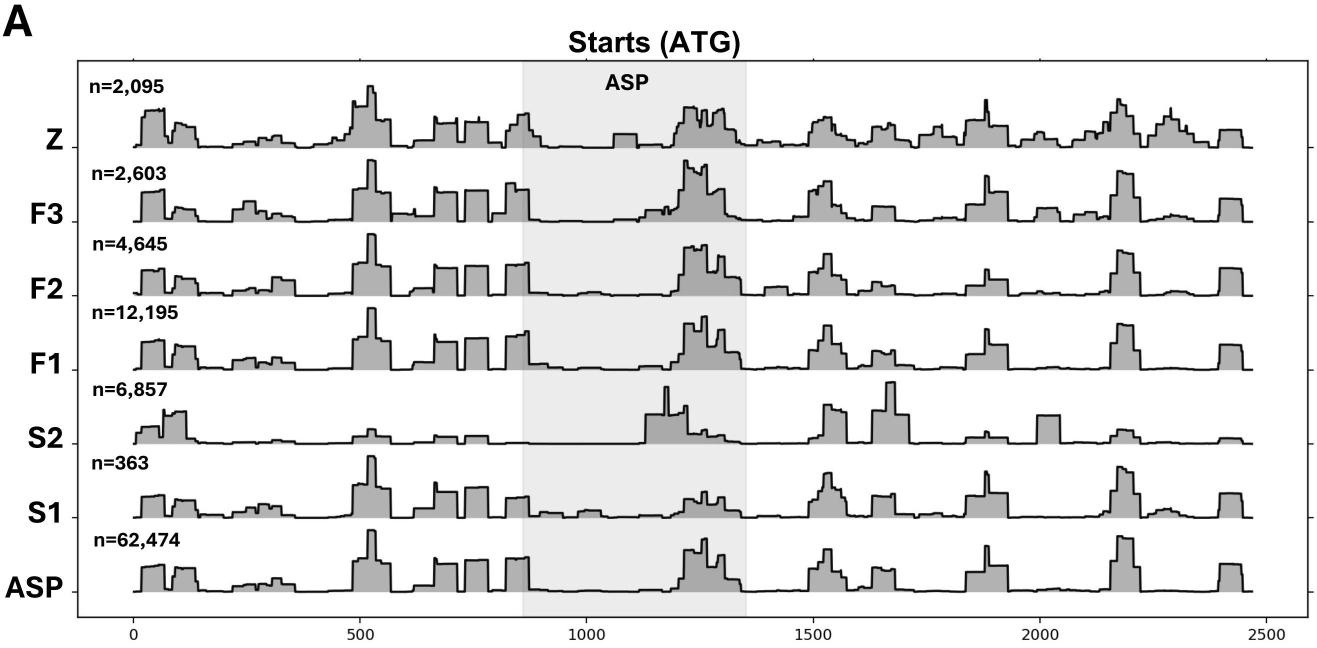


**B**

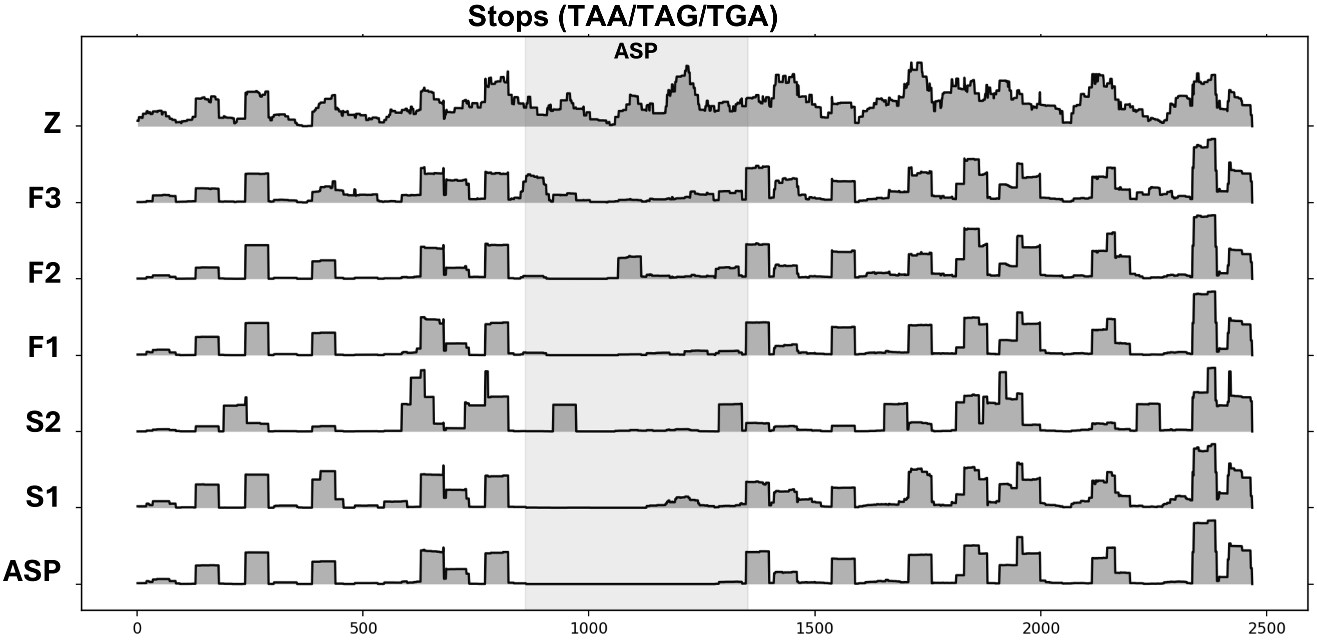


**Supplementary Fig. S6. Ridgeline plots of the start and stop-codon counts in the *env* −2 frame. A** Start-codon (ATG) and **B** stop-codon (TAA/TAG/TGA) densities are shown as ridgeline plots for each type in the 3′ → 5′ direction. The *x*-axis represents alignment coordinates. The *y*-axis represents the counts normalized per sequence at each column (−2 frame only). The shaded band marks the *asp* region.

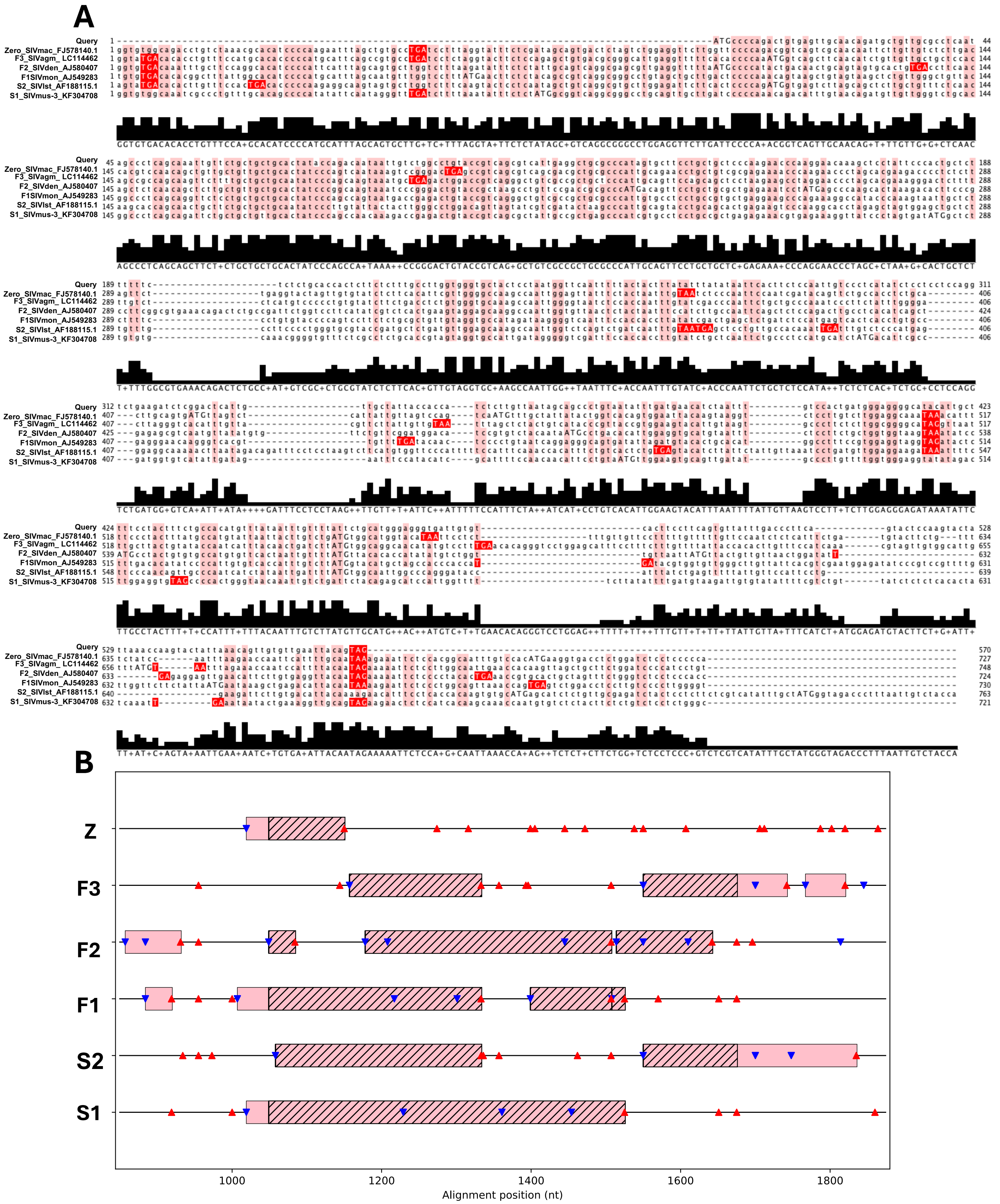

**Supplementary Fig. S7. Multiple sequence alignment and −2 frame stop-codon/ORF visualization for *asp* Evolutionary Group 1. A** Nucleotide multiple sequence alignment of representative Evolutionary Group 1 sequences. Positions sharing ≥80% identity with the *asp* query are highlighted in pink, and in-frame stop codons (TAA/TAG/TGA in −2 frame) are highlighted in red. **B** ORF map of the −2 frame across classes. Horizontal lines show the individual sequences, solid pink rectangles denote the predicted −2 frame ORFs, and hatched segments mark regions overlapping the query-defined window. Blue ▼ indicates start codons and red ▲ indicates stop codons. Positions are in alignment coordinates.

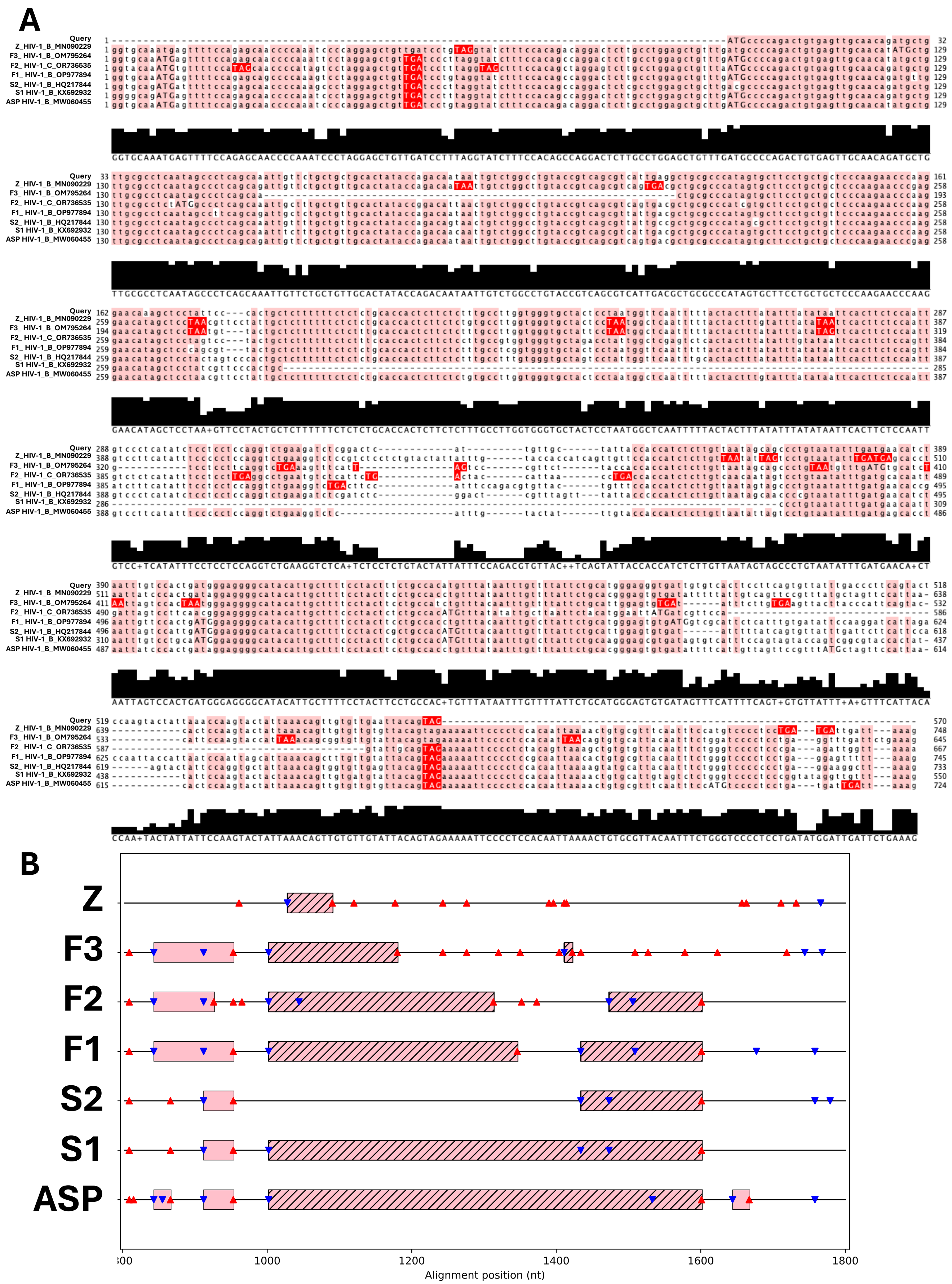

**Supplementary Fig. S8. Multiple sequence alignment and −2 frame stop-codon/ORF visualization for *asp* Evolutionary Group 2. A** Nucleotide multiple sequence alignment of representative Evolutionary Group 2 sequences. Positions sharing ≥80% identity with the *asp* query are highlighted in pink, and the in-frame stop codons (TAA/TAG/TGA in −2 frame) are highlighted in red. **B** ORF map of the −2 frame across classes. Horizontal lines show the individual sequences, solid pink rectangles denote the predicted −2 frame ORFs, and hatched segments mark the regions overlapping the query-defined window. Blue ▼ indicates start codons and red ▲ indicates stop codons. Positions are in alignment coordinates.

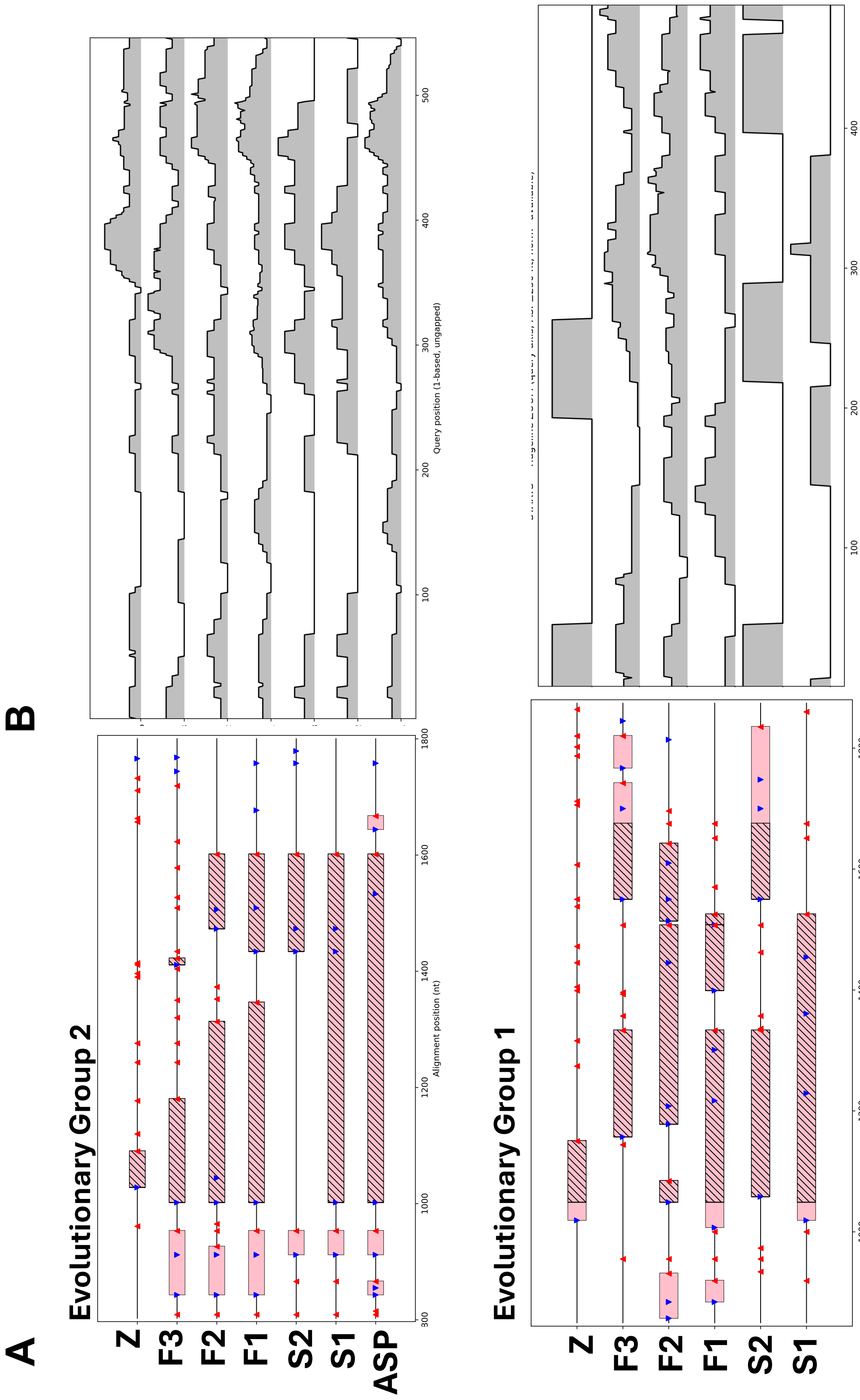


**Supplementary Fig. S9. −2 Frame ORF mapping and start-codon ridgeline plots across the *asp* region in Evolutionary Groups 1 and 2. A** schematic ORF maps of representative sequences from Evolutionary Groups 1 and 2 across the *asp* region. Horizontal lines represent the individual sequences, solid pink rectangles denote the predicted −2 frame ORFs, and hatched segments indicate the overlap with the query-defined window. Blue ▼ indicates start codons and red ▲ indicates stop codons. Positions are shown in alignment coordinates, spanning approximately 800–1800 nt (the *asp* region ±200 nt flank). **B** ridgeline plots showing the per-type start-codon (ATG) density across the *asp* region, normalized per sequence. Positions are in query coordinates (±200 nt around the *asp* region).

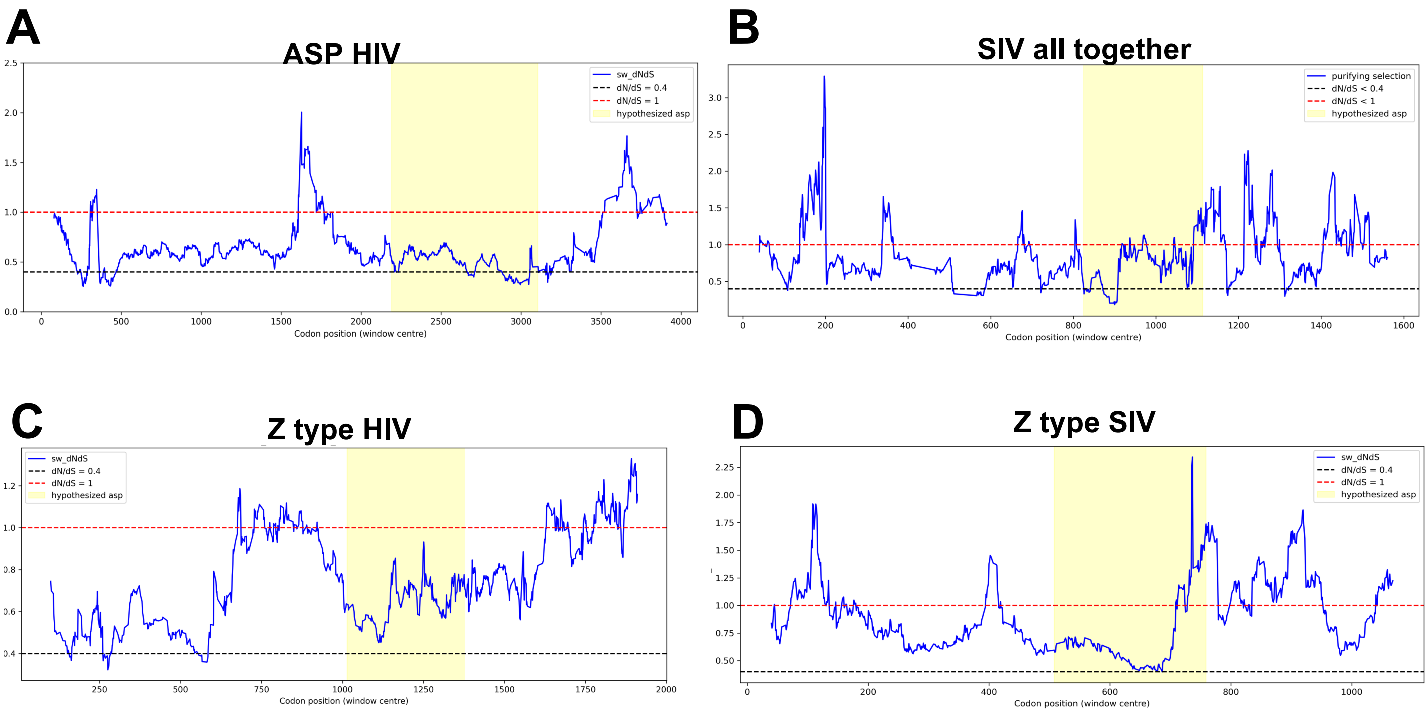


**Supplementary Fig. S10. Sliding-window dN/dS across the *asp*-overlapping region in HIV-1 and SIV.** Sliding-window dN/dS profiles are shown: **A**, HIV-1 ASP type sequences; **B**, SIV sequences combined together; **C**, HIV-1 Z type sequences; and **D**, SIV Z type sequences. dN is the rate of nonsynonymous substitutions, whereas dS is the rate of synonymous substitutions. Accordingly, dN/dS values of <1 indicate purifying selection, values near 1 indicate neutral evolution, and values of >1 may indicate positive selection or relaxed constraint. The blue line shows the dN/dS values across codon-based sliding windows and the yellow band marks the hypothesized *asp*-overlapping region in alignment coordinates. Red dashed reference lines indicate dN/dS = 1.0, representing neutral evolution, and grey dashes dN/dS = 0.4, representing strong purifying constraint.

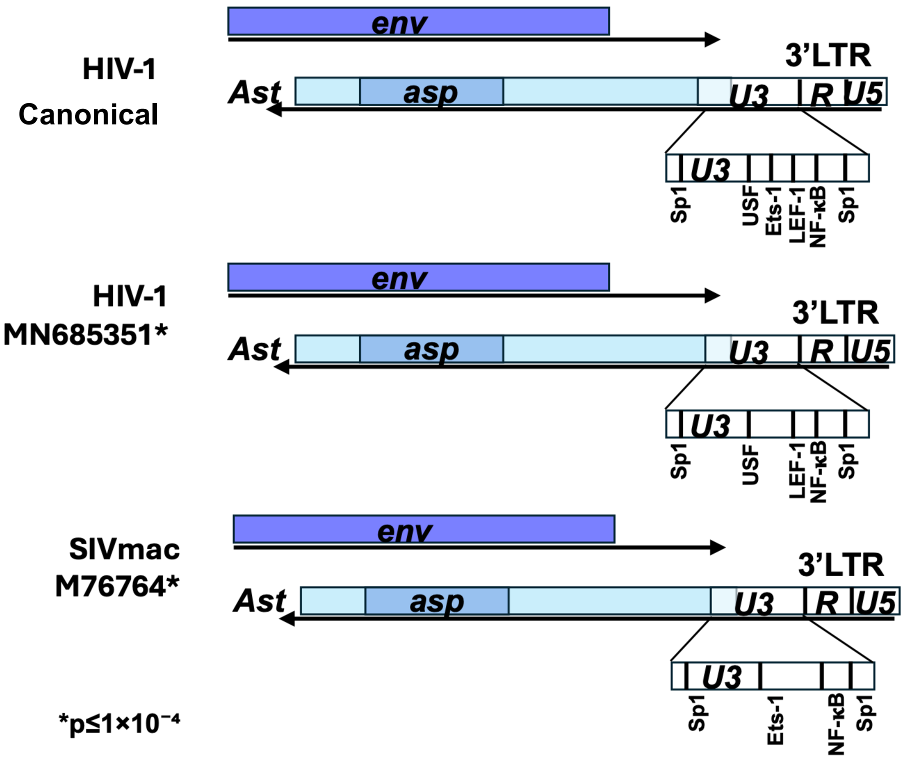


**Supplementary Fig. S11. Representative antisense promoter-associated TF motifs in HIV and SIV U3/LTR-like regions.** Schematic comparison of the canonical HIV-1 antisense promoter TF motif organization with representative HIV-1 (MN685351) and SIVmac (M76764) 3′-U3/LTR-like regions. The top schematic shows the typical arrangement of the TF binding motifs previously associated with HIV-1 antisense promoter activity. Representative HIV-1 and SIVmac LTR-like regions were extracted using GenBank LTR-related annotations or 3′-R-repeat landmarks and scanned with FIMO. Motifs shown for the representative sequences were retained at the stricter threshold of $p \leq 1 \times 10^{-4}$.

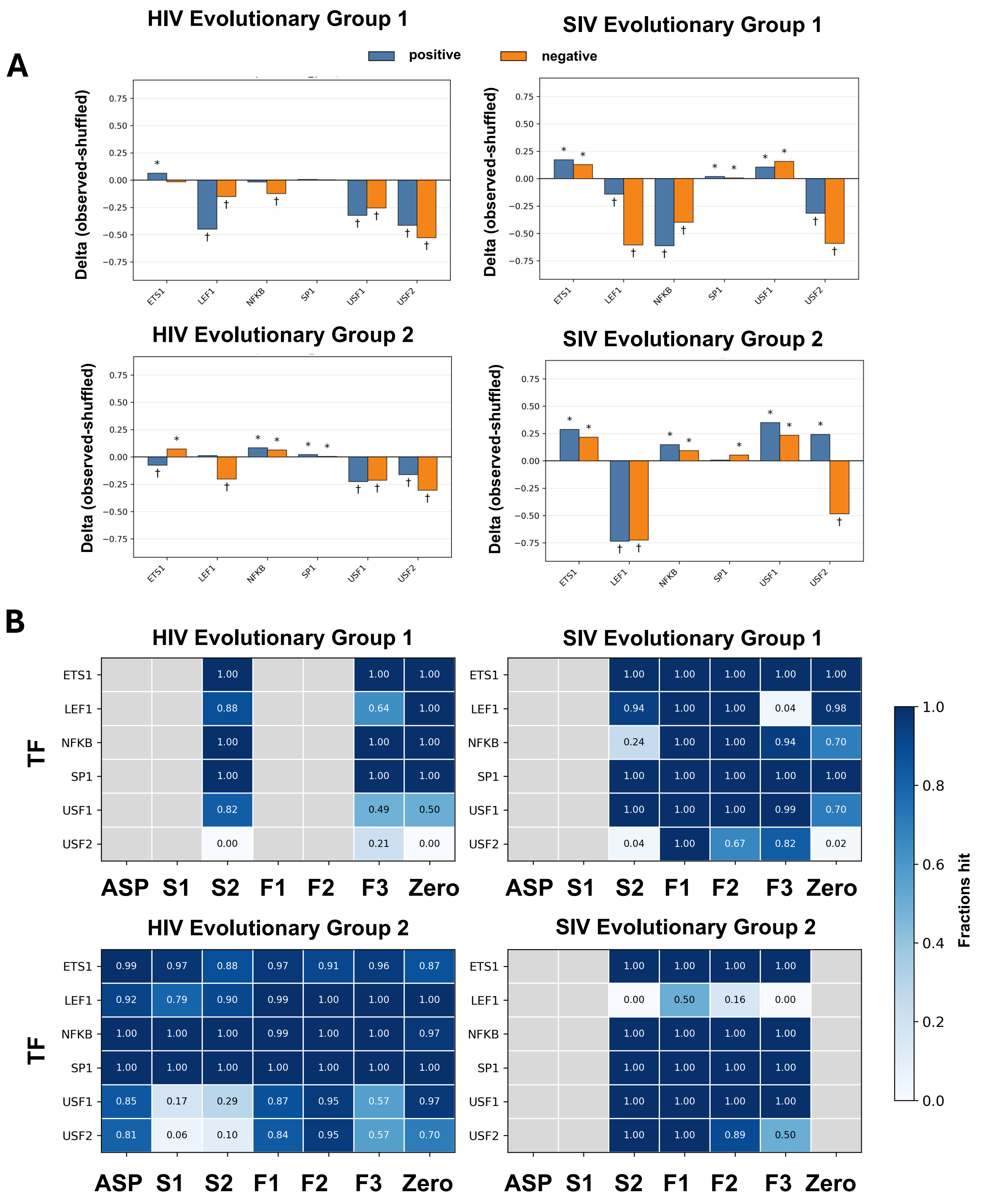


**Supplementary Fig. S12. Distribution of predicted antisense TF promoter-associated motifs across HIV and SIV evolutionary groups at $p \leq 1 \times 10^{-3}$. A** Grouped bar plots show strand-specific enrichment of each TF motif on the positive (+) and negative (−) strands, relative to a dinucleotide-shuffled sequence background (100 replicates per sequence). Each bar is one TF motif. The bar height is the difference between the observed and shuffled-background fraction hit (Δ = fraction of sequences with ≥1 FIMO hit, observed minus background), pooled across ASP types within each virus and evolutionary group. Positive bars indicate enrichment relative to the control and negative bars indicate depletion. **B** Heatmaps of the six TF motifs broken down by individual ASP type. Each cell depicts the fraction of sequences with ≥1 FIMO hits per TF motif (rows) in one ASP type. The panels are separated by virus and evolutionary group. Gray cells indicate ASP types with no available sequences in that category. In **A**, * denotes significant enrichment and † denotes significant depletion relative to the shuffled background (empirical $p < 0.05$, based on the distribution of 100 dinucleotide-shuffled replicates). Bars without a marker did not reach significance in either direction.
